# Engineering NIR probes to enhance affinity and clinical workflow compatibility for prostate cancer imaging

**DOI:** 10.1101/2025.09.16.676549

**Authors:** Gauri S. Malankar, Dani A. Szafran, Gourav Kumar, Joshua Pace, Mackenzie Devereux, Kai Tao, Michelle Gomes, William S. Greer, Cody C. Rounds, Anas M. Masillati, Seseel Gergis, Hayden Ledvina, Melissa H. Wong, Mark J. Niedre, Lei G. Wang, Summer L. Gibbs

## Abstract

Positive surgical margins following radical prostatectomy increase the risk of biochemical recurrence and subsequent disease progression. Fluorescence guided surgery (FGS) using targeted contrast agents has shown clinical benefits for several cancer types. However, current prostate cancer targeted imaging probes exhibit long pharmacokinetic (PK) profiles, necessitating extended waiting periods or repeated hospital visits, limiting their integration into standard clinical workflow. To overcome this critical clinical compatibility challenge, we developed an innovative tri-compartment, chemistry-driven probe design strategy. Specifically, we developed a congeneric library of near infrared (NIR) water soluble fluorescent probes incorporating: (1) a glutamic acid-urea-lysine (EuK) ligand targeting prostate specific membrane antigen (PSMA); (2) a NIR heptamethine cyanine fluorophore optimized for enhanced PSMA binding via secondary binding sites interactions; and (3) distinct PK modulators residing outside the PSMA binding pocket to promote rapid off-target tissue clearance. While molecular docking scores, photophysical properties and live-cell staining results showed similar overall performance, probes bearing PK modulators produced stronger tumor-specific fluorescence in vivo than the control lacking a PK modulator. This effort enabled identification of a lead probe with robust tumor targeting and accelerated off-target clearance, providing optimal tumor-specific signal and contrast in a timeframe, fully compatible with robotic-assisted radical prostatectomy (RARP) timelines.

## Introduction

Prostate cancer is the second most diagnosed cancer in men and the second leading cause of cancer death.^[1]^ For treating patients with localized and locally advanced prostate cancer (PCa), the two primary curative approaches are radical prostatectomy (RP) and radical radiotherapy (RT). However, both treatments carry significant risks of side effects, including urinary incontinence, bowel issues, and erectile dysfunction, all of which can severely impact patients’ quality of life. ^[2] [3] [4] [5] [6] [7] [3] [8] [9]^ Nerve-sparing, robotic-assisted radical prostatectomy (RARP) was developed to reduce postoperative urinary incontinence and erectile dysfunction by preserving the neurovascular bundles adjacent to the prostate. ^[6] [10]^ However, the proximity of these bundles to the prostate capsule can increase the risk of positive surgical margins, which remain a significant challenge even for experience surgeons. A key concern for surgeons is the occurrence of positive surgical margins, which are reported to vary widely, with incidence up to 60% depending on the surgical center and surgeon experience. ^[11] [12] [13]^ Achieving complete PCa resection is particularly difficult without a reliable intraoperative method to identify microscopic tumor deposits at the margin.^[14]^ Macroscopic visual differentiation between cancerous and normal tissue is complex, as little visual white light acuity exists to differentiate benign from malignant tissues, leading to incomplete resections or unnecessary removal of health tissues resulting in postoperative morbidity (i.e., incontinence, impotence).^[15]^ The presence of positive surgical margins is strongly correlated with a higher risk of disease progression and reduced disease-free survival.^[16]^ Notably, patients with positive surgical margins face a two-to five-fold increased risk of biochemical recurrence and increased likelihood of cancer metastases compared to those with negative margins.^[14] [17] [12]^ Thus, development of an intraoperative tool for real-time detection and accurate localization of cancerous lesions could enable surgeons to identify and visualize prostate cancer intraoperatively, facilitating complete cancerous tissue resection while preserving other critical structures, reducing the risk of recurrence and potentially improving long-term outcomes for patients.

Prostate-specific membrane antigen (PSMA), also known as glutamate carboxypeptidase II, is a well-established prostate cancer biomarker. ^[18] [19] [20] [21]^ The PSMA protein is highly overexpressed in prostate cancers, and membrane bound in the extracellular space, making it a desirable target for detection and treatment of prostate cancer. ^[22]^ To date, various PSMA targeted imaging agents have been developed, including fluorescent PSMA-activatable probes, ^[23] [24] [25] [26] [27] [28] [29] [30]^ PSMA-antibody and minibody based probes, ^[31] [32] [33]^ peptide-based probes, ^[34]^ hybrid tracers, ^[31]^ radiolabeled small-molecule probes,^[35] [36]^ and radiolabeled monoclonal antibodies.^[37]^ Among these PSMA targeting ligands, glutamate-urea-lysine (i.e., EuK), a urea-containing small molecule, has previously been developed for prostate cancer specific targeting. EuK is uniquely positioned as an optimal PSMA targeting ligand for RARP, where a well-defined pharmacokinetic (PK) profile is critical for compatibility with the existing surgical workflow. The clinical utility of this ligand is further demonstrated by clinically approved imaging agents, such as ^18^F-DCFPyl and ^68^Ga-PSMA-617, both based on EuK PSMA targeting, which are currently utilized for prostate cancer diagnosis and staging. ^[38] [39] [40]^ However, these radiolabeled EuK derivatives cannot be readily imaged in the operating room and thus are not suitable as an intraoperative aid.^[41]^ Secondary to tumor resection during RARP, the small delicate cavernous nerve, which is responsible for continence and potency, is often damaged, with reported incidence rate as high as 60%.^[42]^ Patients thus frequently suffer from postoperative complications due to both inadequate tumor control and iatrogenic nerve injuries, highlighting the critical need for an improved intraoperative visualization technique.

Fluorescence-guided surgery (FGS) offers the potential for improved visualization of specifically highlighted tissues (i.e., cancer, nerves, blood vessels) intraoperatively in real time. ^[43]^ Using optical imaging technology, FGS enables real time, minimally invasive visualization through judiciously designed imaging agents coupled with surgical imaging systems that can integrate into current surgical workflows. FGS using tumor-specific tracers is a promising technique to highlight cancer cells and facilitate real-time identification, visualization and resection of cancerous tissues or preservation of critical structures, such as nerves or vasculature.^[44]^ Clinically, intraoperative imaging systems operating nearly exclusively in the near infrared (NIR, 650-900 nm) spectral window are employed successfully for FGS, with fluorescent contrast agents administered either intravenously or orally (i.e., methylene blue, indocyanine green, aminolevulinic acid/protoporphyrin IX).^[45] [46] [43] [47]^ Recently, two NIR molecularly targeted probes (i.e., pafolacianine and pegulicianine) have received FDA approval for intraoperative cancer detection across major indications (ovarian, lung, breast). ^[48]^ However, a FGS imaging agent specifically optimized for prostate cancer resection and compatible with the surgical workflow timelines (i.e., tumor specific contrast within 2-4 hours post-injection) remains an unmet clinical need. An imaging probe that could be administered preoperatively or at the beginning of surgical procedure to provide robust tumor-specific contrast throughout the entire RARP (i.e., within 4-hour window after administration) would offer tremendous clinical benefits by improving surgical precision, preserving vital structures and improving patient outcomes. Notably, >80% of prostatectomies are completed using da Vinci surgical robot (Intuitive Surgical), which is equipped standard with a NIR fluorescence imaging channel compatible with indocyanine green imaging.^[49]^

Current prostate cancer imaging probes under development for FGS (i.e., IS-002 and OTL-78) typically exhibit prolonged pharmacokinetics (PK), which require extended time between injection and surgery to assess cancer location or residual tumor detection. Moreover, slow elimination from non-target tissues can lead to persistent background signal, reduced contrast and delays that interfere with surgical planning.^[17]^ This prolonged imaging process may hinder integration into the current surgical workflow of RARP, challenging clinical adoption. The chemical modifications that influence biodistribution and clearance, such as the introduction of hydrophilic substituents, cationic groups, anionic groups, and pegylation, can be strategically incorporated into the probe structure to optimize their PK profile for enhance clinical suitability. Additionally, tailoring the physicochemical properties of the fluorophore scaffold can simultaneously enhance biomarker binding affinity. Collectively, these structural modifications would enable rapid clearance from healthy tissues while promoting selective and sustained retention in tumor regions. This chemistry-driven PK modulation could significantly enhance the clinical applicability of imaging agents, enabling precise, real-time surgical decision-making during prostate cancer resection.

To address these unmet clinical needs, we developed an innovative tri-compartment FGS probe strategy designed specifically to target prostatic tumors intraoperatively in real-time while maintaining compatibility with current surgical workflows. The lead probe identified through this effort showed rapid extravasation, efficient tumor perfusion, sustained tumor retention and accelerated background clearance, providing high tumor-specific fluorescence signal and optimal contrast within the critical intraoperative timeframe. Importantly, these probes fluoresce in the 800 nm channel, compatible with clinically established indocyanine green imaging channel widely used during RARP. This clinically integrated approach holds substantial promise to enhance surgical precision, improve oncological outcomes, and could significantly reduce postoperative complications following prostatectomy.

## Results and Discussion

### Tri-Compartment PSMA-targeted Probe Design for Optimal FGS of Prostate Cancer

To overcome the limitations of existing PSMA-targeted intraoperative imaging probes for prostatectomy, we designed a novel tri-compartment PSMA-targeted probe strategy specifically optimized for prostate cancer FGS (Fig.1a). Our design strategy integrated three distinct but complementary chemical features: (1) a small-molecule based EuK ligand to ensure high affinity targeting to prostate cancer protein biomarker, PSMA, by coordinating with the zinc active site of the PSMA protein;^[49] [50]^ (2) a NIR heptamethine cyanine fluorophore tagged with an optimal spacer and anionic sulfonate group designed to engage secondary binding interactions, including hydrogen bonding and amphiphilic interactions within PSMA’s amphipathic entrance funnel to the zinc active site, further enhancing probe-protein target binding affinity; and (3) PK modulators strategically introduced onto the fluorophore’s second indole ring, residing outside the amphipathic entrance funnel to fine-tune extravasation kinetics, tumor tissue binding and off-target tissue clearance, ensuring rapid achievement of maximum tumor-specific contrast within the clinically relevant intraoperative window. Combining these judiciously engineered chemical compartments resulted in a congeneric library of fluorescent probes, creating a uniquely modular and clinically compatible platform designed to identify optimal fluorescent probes for prostate cancer FGS.

**Figure 1.**
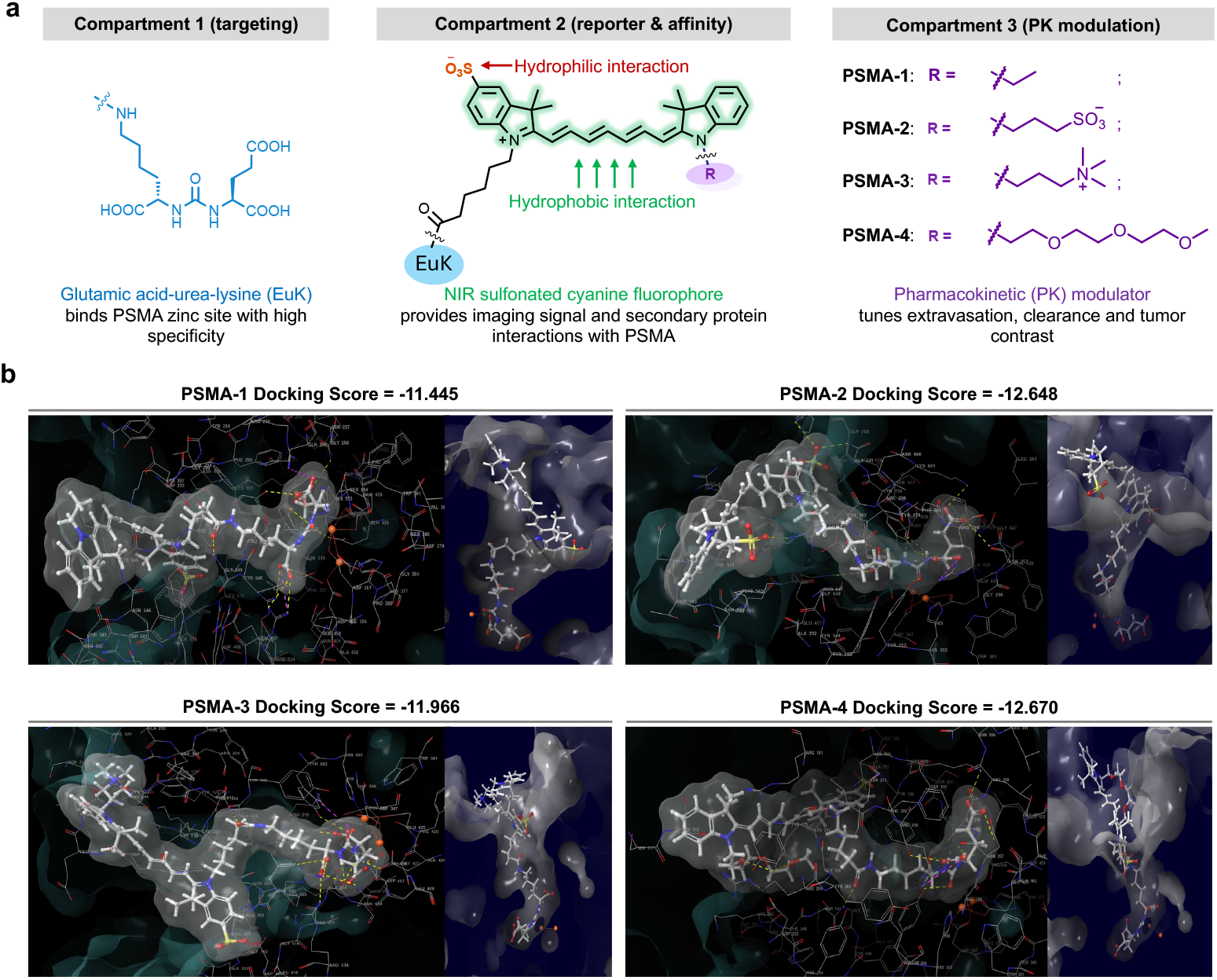
Design strategy for development of NIR, water-soluble PSMA-targeted probes with varied pharmacokinetic (PK) modulators. **a.** Schematic illustration of the tri-compartment PSMA-targeted probe design strategy. Compartment 1: PSMA binding moiety EuK (glutamic acid-urea-lysine), which coordinates with the zinc active site in the PSMA protein. The EuK binding moiety is tagged with a linker of optimal length to position the ligand into the active site while the fluorophore remains on the protein surface. Compartment 2: NIR hydrophobic heptamethine cyanine dye selected for fluorescence compatibility with the NIR imaging channel in clinically approved FGS systems (e.g., da Vinci Surgical Robot). The heptamethine cyanine fluorophore contains a hydrophilic anionic sulfonate group designed to promote secondary binding interactions, enhancing probe binding affinity of the PSMA protein. Compartment 3: pharmacokinetic (PK) profile modulators incorporated onto the fluorophore with the position opposite from the EuK ligand, designed to facilitate rapid off-target clearance. **b**. Molecular docking analysis of PSMA probes in human glutamate carboxypeptidase II (i.e., PSMA). Molecular docking studies were performed using Schrödinger Maestro interface to define the binding modes and docking scores. The surface representation from the top view showed the position of each PSMA probe within the PSMA protein active site cavity. The zinc ions are shown as orange spheres and the protein is represented as a gray cartoon.

### Molecular Docking Analysis of Protein-Ligand Interactions Enabled Rational PSMA-targeted Probe Design

The optimal spacer length between the fluorophore and EuK ligand as well as the role of the fluorophore as a secondary binding motif to enhance the overall PSMA binding affinity were evaluated using molecular docking studies for each of the designed probes against the X-ray crystal structure of PSMA (PDB code 2XEG) ^[50]^ to elucidate each designed probe’s potential binding modes within the active site of PSMA (Fig. 1b). Docking studies revealed that the EuK moiety with the spacer occupied a buried position in the active site of the PSMA protein, while the cyanine fluorophore was localized onto the surface of the protein. In all the PSMA structures, the carboxylate group of the EuK ligand participated in hydrogen-bonding interactions with ARG534 and ARG536 in the arginine patch (S1 subpocket), while the carbonyl carbon of the uriedo formed a hydrogen bond with TYR552. Furthermore, the beta and gamma carboxylate groups of EuK formed electrostatic interactions with ARG210 and LYS699 residues of S1’ pharmacophore subpocket. The beta carboxylic acid moiety of the EuK engaged hydrogen bonding with TYR700 and the gamma carboxylate group similarly formed a hydrogen bond with ASN257. The uriedo nitrogen atoms formed hydrogen bonds with GLH424 and the GLY518 main chain carbonyl in the S1’ pocket, and the carbonyl oxygen directly interacted with the catalytic zinc atom Zn1751. Notably, the sulfonate group on the cyanine fluorophore formed hydrogen-bonding interactions with GLY702 within the S1’ pocket, consistent with previous findings. ^[51]^ The carbonyl group of the linker also formed a hydrogen bond with GLY548 in the S1’ pocket. Importantly, the PK modulators in PSMA-2 and PSMA-4 (the sulfonate and PEG containing compounds, respectively) further contributed hydrogen bonding interactions with LYS207 in the S1’ pocket, illustrating the multifunctional role of these engineered structural motifs in enhancing probe-protein target binding affinity.

### Synthesis of Engineered PSMA-targeted Fluorescent Probes

An overview of the synthetic scheme of PSMA-targeted fluorescent probes is provided as follows (Fig. 2a). The indole ring **3** was synthesized via Fisher-indole cyclization by combining sulfonate substituted hydrazine with 3-methyl-2-butanone under acidic conditions. The formed indole derivatives were converted to potassium salt in the presence of potassium hydroxide and isopropanol following a modified literature protocol.^[53]^ The crude product was washed with diethyl ether and vacuum dried overnight to give compound **3**, which was subsequently condensed with commercially available linker **4** through an N-alkylation reaction to form compound **5**. The additional indole derivatives (**7**-**10**) were synthesized by N-alkylation reactions with alkylation agents. These N-alkylated indole derivatives were purified by recrystallization procedures. The open-chain heptamethine cyanine fluorophores were synthesized by condensing compound **5** with the commercially available linker **6** in the presence of acetic anhydride to form the imine intermediate. The indole derivatives (**7**-**10**) were subsequently added under basic conditions to afford asymmetric cyanine dyes as the major product (**11**-**14**). In the next step, the fluorophore was conjugated to ligand **15**, followed by removal of the t-butyl group under acidic conditions. The final EuK conjugated fluorescent probes were purified by HPLC and obtained as green solid compounds (**16**-**19**) in 20-30% yield. Probe synthesis and characterization details are provided in the Supplementary Information.

**Figure 2.**
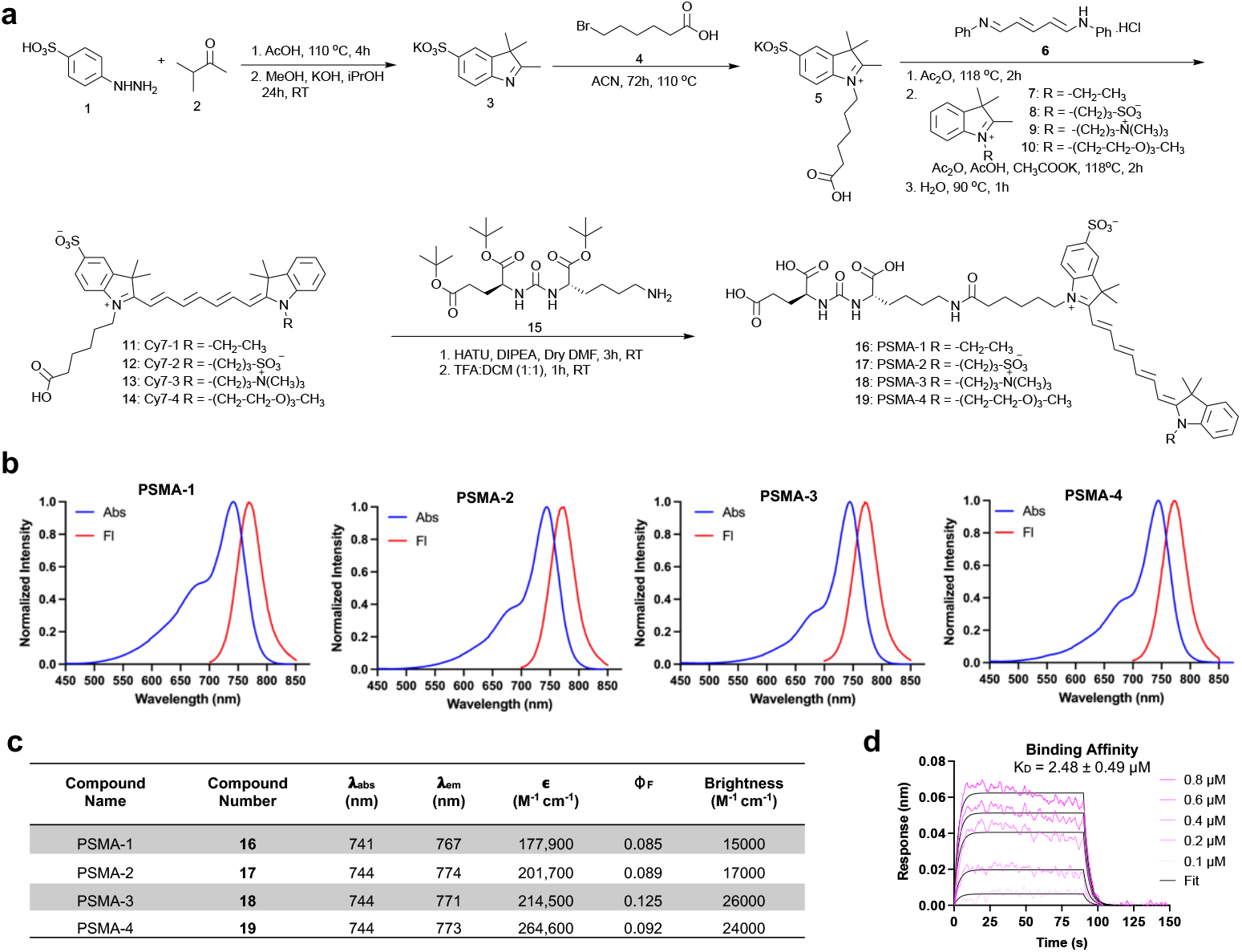
Synthetic route, photophysical property and target binding of NIR PSMA-targeted probes. **a.** Synthetic scheme for preparing NIR PSMA-targeted probes PSMA-1 (**16**), PSMA-2 (**17**), PSMA-3 (**18**), and PSMA-4 (**19**). **b**. Normalized absorption and emission spectra of the synthesized PSMA-targeted probes collected at a concentration of 10 µM in phosphate buffer saline (PBS, pH 7.4) containing 1% dimethyl sulfoxide (DMSO). The displayed spectra represent the average of n = 3 spectra normalized to their respective maxima. Abs = Absorbance (blue), Fl = Fluorescence (red). **c**. Tabulated spectral properties of PSMA-targeted probes in aqueous media. Λ_abs_ = maximum absorbance wavelength, λ_em_ = maximum emission wavelength, ε = extinction coefficient at λ_abs_, Φ_F_ = fluorescence quantum yield using HITCI (1,10,3,3,30,30-hexamethylindotricarbocyanine iodide) as a reference standard. **d**. The target binding of the PSMA probes was measured by Biolayer Interferometry (BLI). Representative sensograms show PSMA-4 binding to recombinant PSMA immobilized on streptavidin (SA) biosensors. Association was measured at probe concentrations from 0.1-0.8 μM, followed by dissociation in buffer without analyte. Experimental data (colored curves) were globally fit to Langmuir 1:1 binding model (black curves) using ForteBio Data Analysis HT 10.0 to obtain association rate (k_a_), dissociation rate (k_d_), and the equilibrium dissociation constant (K_D_). The data were acquired using ForteBio Octet^®^RED384 instrument with Acquisition software 9.0.

### Photophysical Property Characterization of PSMA-targeted Fluorescent Probes

The photophysical properties of all the synthesized PSMA-targeted probes were characterized by measuring absorption and emission spectra in a phosphate-buffered saline (PBS, pH 7.4) solution containing 1% dimethyl sulfoxide (Fig. 2b). All probes showed spectral compatibility with clinical FGS systems, with absorption maxima ranging from 741 to 744 nm and emission maxima ranging from 767 to 774 nm, yielding a Stokes shift of ∼30 nm. Molar extinction coefficient ranged from 177,900 to 264,600 cm^−1^ M^−1^ for PSMA-targeted probes in PBS. When comparing the quantum yield of these probes, PSMA-3 has the highest quantum yield among the series at 12.5%. The calculated brightness for the probes ranged from 15,000 to 26,000 cm^−1^ M^−1^ (Fig. 2c). These data collectively suggested that the introduction of the various PK modulators minimally impacted the spectral and photophysical properties of the cyanine fluorophore label.

### Binding Affinity Quantification of PSMA-targeted Probes using Biolayer Interferometry

A biolayer interferometry (BLI) assay was used to experimentally confirm the computational docking results and evaluate the binding affinities of the synthesized PSMA-targeted probes. Binding kinetics were measured using biotinylated recombinant PSMA protein (extracellular domain K44-A707) immobilized on streptavidin (SA) biosensors with a calculated average association rate constant k_a_ = (2.48 ± 0.39) x 10^5^ M^-1^s^-1^, an average dissociation rate constant k_d_ = (3.1 ± 0.1) x 10^-1^ s^-1^ and an average equilibrium dissociation constant K_D_ = 2.48 ± 0.49 µM (Fig. 2d). These BLI finding further supported the probe design strategy for PSMA targeting, establishing the molecular basis for subsequent *in vitro* and *in vivo* characterization studies aimed at optimizing PSMA-targeted fluorescent probes for RARP FGS.

### *In vitro* PSMA Specificity Characterization in Prostate Cancer Cell Lines

Live-cell staining studies on prostate cancer cell lines with varied PSMA expression levels to characterize PSMA probe specificity was completed. PSMA expression was confirmed by immunofluorescence staining using the PSMA targeted antibody, Mouse Hybridoma Prost J533, directly conjugated to Alexa Fluor 488 in C4-2 (high), 22RV1 (moderate), and PC-3 (negative) prostate cancer cell lines. The live-cell imaging results demonstrated clear correlation between PSMA expression levels and probe fluorescence signals across all three tested prostate cancer cell lines. In particular, the PSMA-positive C4-2 cells showed the highest fluorescence intensity compared to medium PSMA expression level 22RV1 cells, whereas minimal signal intensity was observed in PSMA-negative PC-3 cells, confirming the specificity of our PSMA-targeted probes (Fig. 3a). Quantitative analysis of single-cell fluorescence signal further confirmed this correlation with C4-2 showing the highest intensity values, followed by medium level in 22RV1 cells and minimal signal in PC-3 cells (Fig. 3b). As a control group, a fluorophore without the EuK ligand (**14**) didn’t show any fluorescence signal in any of the cell lines tested (Fig. S2). These *in vitro* live-cell imaging results confirm that all the probes specifically target the PSMA protein in prostate cancer cell lines, validating the expected target specificity and indicating variations in the indole ring substitution with PK modulators had minimal impact on *in vitro* cellular PSMA specificity.

**Figure 3.**
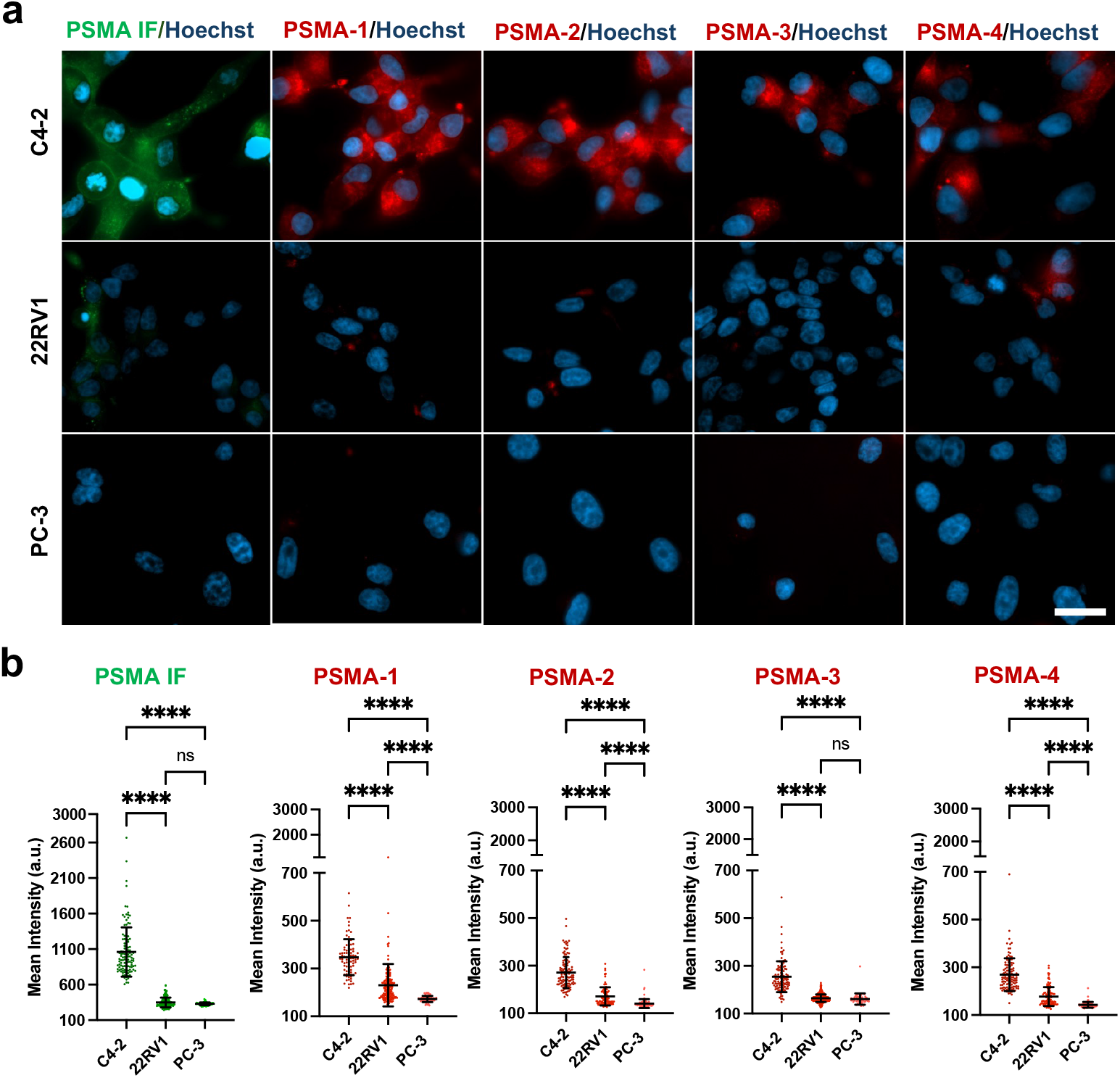
In vitro live-cell imaging of PSMA-positive and PSMA-negative cells by fluorescence microscopy. **a.** PSMA expression levels of C4-2 (PSMA++), 22RV1 (PSMA+), and PC-3 (PSMA-) prostate cancer cells were confirmed by PSMA immunofluorescence (IF) staining with PSMA antibody (Mouse Hybridoma Prost J533) directly conjugated to AF488 (left panel, green). Subsequent cell staining with PSMA-targeted probes, PSMA-1, PSMA-2, PSMA-3 and PSMA-4 was completed with each image normalized to intensities observed in the C4-2 cell line of the respective probe (red). Cell nuclei were stained with Hoechst 33342 (blue). Scale bar = 20 μm. **b**. Quantitative image analysis of the fluorescence intensities of these costained samples showed the expected PSMA expression pattern with IF as well as PSMA probe staining. Significant differences in mean fluorescence intensity are marked with asterisks between prostate cancer cell lines, where significance was evaluated using one-way ANOVA, and ^**^ = p value <0.01, ^***^ = p value <0.001, ^****^ = p value <0.0001.

### Noninvasive Blood PK Assessment of PSMA-targeted Probes by Measurement of Diffuse *in vivo* Fluorescence

Noninvasive Diffuse in vivo fluorescence imaging was utilized to quantify the impact of PK modulators on PSMA probe circulation dynamics *in vivo*. We performed in vivo measurements of PSMA probe fluorescence using our previously described noninvasive diffuse in vivo flow cytometry (DiFC) instrument. DiFC enables real-time, noninvasive monitoring of intensity changes in live mice using diffuse fluorescence detection, eliminating the need for serial blood sampling or animal euthanasia (Fig. 4a).^[52] [53]^ Briefly, DiFC measurements were performed by placing the optical fiber probe on the skin over the vessels with ultrasound gel applied for improved light delivery and collection (Fig. 4b). The instrument photomultiplier tube (PMT) gain was adjusted such that the signal coming from a vehicle-injected control mouse was recorded as baseline fluorescence, and was then kept at the same value for all subsequent experiments. After baseline fluorescence data was collected for each mouse, 10 nmol of each PSMA-targeted probe was administered via retroorbital injection. Mice were then continuously scanned until the fluorescence measurement returned to the baseline value as quantified prior to probe injection, reflecting complete probe clearance from systemic circulation. The probe clearance kinetics (i.e., extravasation rates) varied distinctly based on the PK modulating motif included in the PSMA probe (Fig. 4c). Specifically, the clearance time of PSMA-1 (without PK modulator) and PSMA-3 (with quaternary amine-based PK modulator) were very similar, each returning to the baseline 4 hours post-injection. In contrast, PSMA-2 (with sulfonate-based PK modulator) showed the slowest clearance rate, requiring ∼10 hours to return to baseline fluorescence levels, while PSMA-4 (with PEG-based PK modulator) demonstrated the fastest blood clearance, returning to baseline fluorescence levels within just 2 hours of systemic administration (Fig. 4c). These blood PK profiles clearly highlight the significant impact of PK modulators on *in vivo* probe extravasation rates and systemic clearance, which was hypothesized to directly impact tumor probe uptake rate, tumor-specific contrast and imaging performance during FGS.

**Figure 4.**
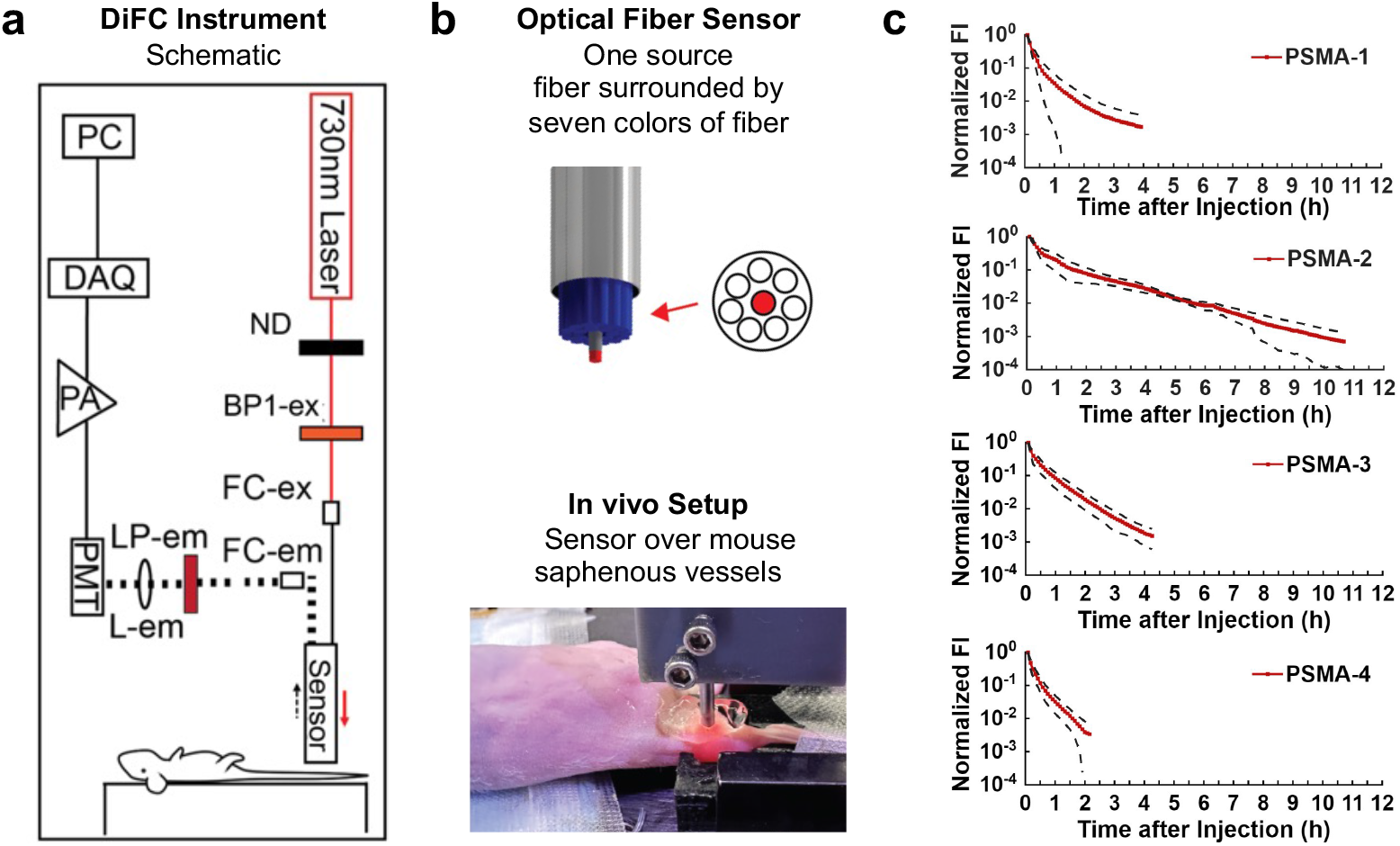
Noninvasive Blood PK Assessment using Diffuse in vivo Fluorescence Measurements. **a.** In the DiFC instrument a 730 nm laser with a 720/24nm excitation filter (BP1-ex) was used for excitation with the light power at the sample tuned to 25 mW using a neutral density (ND) filter. The laser was coupled into a source fiber with a collimation package (FC-ex). The output of the collection fibers was collimated (FC-em) and the light was passed through a 780 nm long pass emission filter (LP-em) before being focused on to the surface of a photomultiplier tube (PMT) with a 30 mm focal length lens (L-em). The output signals from the PMTs were filtered with an electronic 100 Hz low pass filter, amplified with a low-noise voltage pre-amplifier (PA), and then acquired with a data acquisition board (DAQ). **b**. The DiFC instrument utilized a custom-designed optical fiber sensor had one 300 µm excitation fiber surrounded by seven 300 µm collection fibers for fluorescence detection. The DiFC optical fiber sensor was placed on the skin above the saphenous vessels in the mouse hind limb. **c**. The fluorescence signal was continuously monitored after administration of the PSMA probes (PSMA-1, PSMA-2, PSMA-3, and PSMA-4) via retroorbital injection to quantify probe extravasation rates. The dashed lines (black) represent the error bars for each group, and the solid line (red) shows the average of the reading (n=4 mice/probe). PSMA-1 (without PK modulator) and PSMA-3 (with quaternary amine-based PK modulator) showed similar extravasation rates, each returning to the baseline 4 hours post-injection. PSMA-2 (with sulfonate-based PK modulator) showed the slowest clearance, requiring ∼10 hours to return to the baseline level of fluorescence, whereas PSMA-4 (with PEG-based PK modulator) demonstrated the fastest blood clearance, returning to the baseline level of fluorescence within 2 hours. Fl = Fluorescence.

### *In Vivo* Assessment of PK Modulator-Driven PSMA-positive Tumor Targeting

The PSMA-targeted probes were further evaluated in murine models of prostate cancer to assess the impact of the PK modulators on the tumor-specific uptake, retention and contrast as well as off-target clearance *in vivo*. PSMA-positive tumors were subcutaneously xenografted into mice by injecting the C4-2 prostate cancer cells, where tumors were allowed to grow for 6-8 weeks. Once the tumor reached 500-700 mm^3^, the mice were intravenously injected with 10 nmol of each probe. White light and fluorescence images were collected at the clinically relevant 4-h timepoint after systemic administration (Fig. 5a).^[54] [55]^ Fluorescence images were collected using a 740 nm excitation filter and 780 nm longpass emission filter. The PSMA-positive tumors showed significantly higher fluorescence intensity after systemic administration of PSMA-2, PSMA-3 and PSMA-4 as compared to PSMA-1. PSMA-1, a control probe contained an ethyl group instead of a PK modulating group, providing weak tumor-specific fluorescence signal. Additionally, PSMA-2 demonstrated the highest tumor-specific fluorescence intensity as compared to the other three PSMA-targeted probes (Fig. 5b). Furthermore, using muscle as the reference background tissue, the calculated signal-to-background ratios (SBR) also showed that PSMA-2 provided the highest tumor-specific contrast at the 4-h imaging timepoint (Fig. 5c). Notably, despite their similar photophysical properties, docking scores and *in vitro* cell staining profiles, the PK modulators on the fluorophore scaffold not only impacted the probe’s *in vivo* extravasation rates, but also their tumor-specific uptake and off-target tissue clearance kinetics. In particular, the enhanced tumor-specific fluorescence signals observed from PSMA-2 and PSMA-4 probes coincide with additional hydrogen-bonding interactions between their respective PK modulators and the PSMA protein (Fig. 1b), which likely increased tumor-specific accumulation compared to PSMA-1 (control probe without a PK modulating moiety) and PSMA-3 (without hydrogen-bonding interaction). Together, these properties highlight the critical role of PK profile modulations in optimizing targeted FGS probe performance and enhancing clinical translation potential.

**Figure 5.**
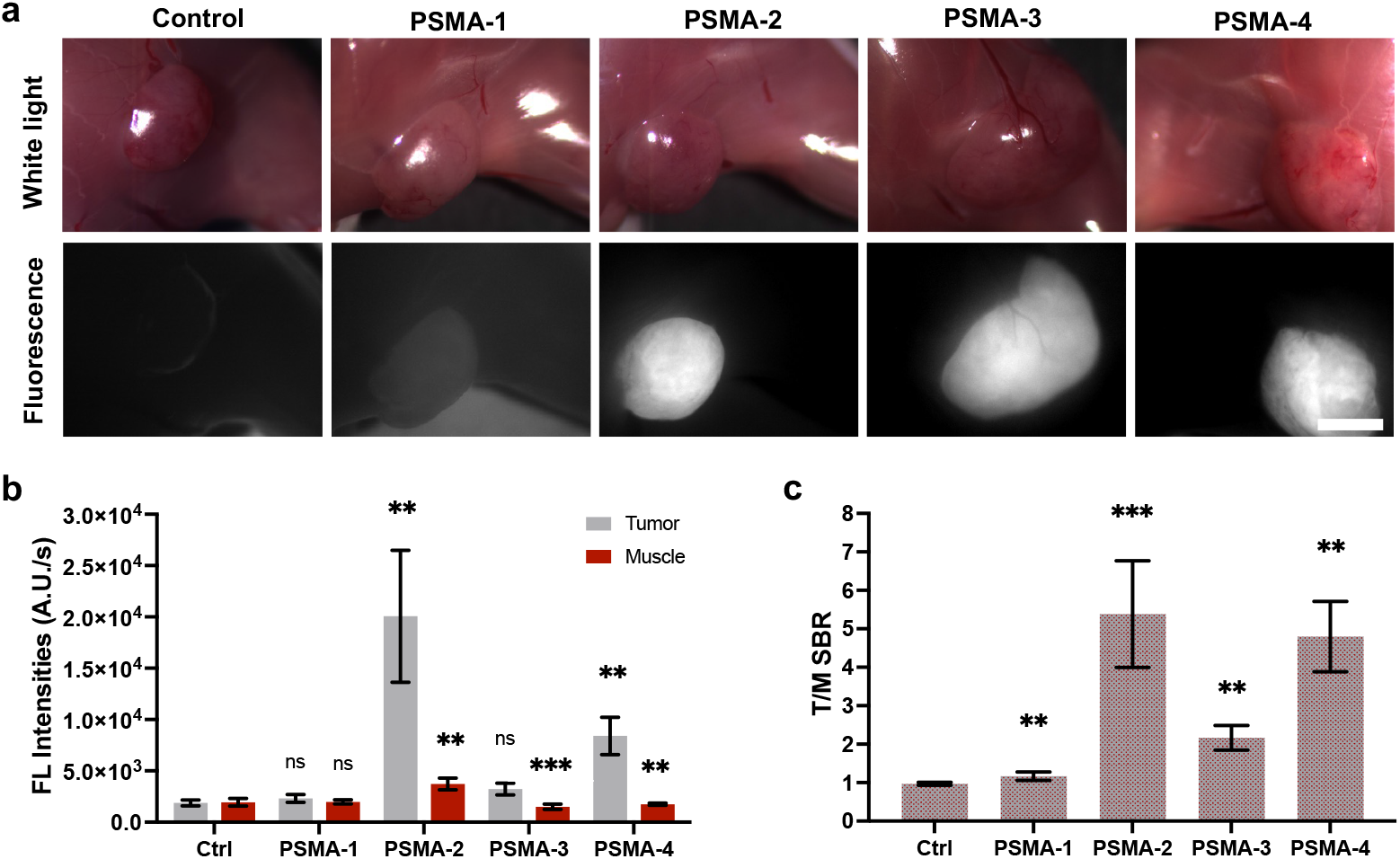
In vivo quantification of PSMA-targeted probes in PSMA-positive tumor-bearing mice. **a.** Representative white light and fluorescence images of PSMA-positive tumor-bearing mice following intravenous injection of PSMA-1, PSMA-2, PSMA-3, or PSMA-4 (10 nmol each). Fluorescence images were acquired 4 hours after probe administration using 740 nm excitation and 780 nm emission filters. Scale bar = 4 mm. **b**. The tissue-specific fluorescence intensities from tumor and muscle in both vehicle-injected control and probe-injected groups (n = 4 mice/group) were quantified enabling **c**. calculation of the tumor-to-muscle signal-to-background ratio (SBR). PSMA-2, containing a sulfonate-based PK modulator, exhibited the highest tumor-specific intensity and SBR among all tested probes, followed by PSMA-4. Ctrl = vehicle-injected control PSMA-positive tumor-bearing mice for autofluorescence quantification; FL = fluorescence. The statistical significance was evaluated using one-way ANOVA with gaussian distribution followed by Fisher’s LSD test, where ^**^ = p value <0.01, ^***^ = p value <0.001, ^****^ = p value <0.0001.

## Conclusion

For FGS clinical translation potential, targeted fluorescent probes must not only achieve high tumor contrast and robust safety profiles, but must also seamlessly accommodate clinical workflow constraints. Herein, a tri-compartment probe design strategy for PSMA-targeted fluorescent probes was successfully optimized specifically for prostate cancer FGS. Our modular chemical design integrated a high affinity PSMA targeting ligand (EuK), an optimized NIR heptamethine cyanine fluorophore bearing an anionic sulfonate for enhanced PSMA targeting via secondary protein binding site interactions and strategically selected PK modulators. Importantly, these PK modulators strongly influenced the probe extravasation kinetics, tumor-specific uptake and off-target tissue clearance, all critical characteristics necessary to ensure rapid and robust tumor-specific fluorescent contrast within the timing constraints of the clinical RARP workflow. Protein-ligand docking studies and BLI assays confirmed that the EuK ligand interacts specifically with the zinc metal site on the PSMA protein, validating the rational probe design strategy. While all synthesized probes showed similar photophysical properties and PSMA specificity in cell lines, the noninvasive DiFC imaging revealed significant differences in blood clearance rates that were directly attributed to the PK modulators. Specifically, PSMA-4 containing a PEG-based PK modulator showed the fastest extravasation rate, while PSMA-2 containing a sulfonate-based PK modulator demonstrated the slowest clearance rate. Interestingly, PSMA-3 contained a quaternary ammonium-based PK modulator, which exhibited minimal impact on probe blood PK profile compared to the control probe PSMA-1, which contained an ethyl group instead of a PK modulator. Subsequent *in vivo* tumor-targeting assessment identified the PSMA-2 probe as the optimal FGS candidate due to its balance of extravasation rate, tumor-specific fluorescence and accelerated off-target tissue clearance, providing exceptional tumor-specific contrast within the clinically relevant 4-h surgical window. These findings demonstrate the rationally engineered PK modulators can significantly enhance the suitability of fluorescent probes for real-time prostate cancer FGS, suggesting the necessity of integrating PK modulation into the broader FGS probe development strategy. Critically, these probes need to demonstrate their ability to accommodate the clinical procedural constraints and seamless integration into the clinical workflow, while providing tumor-specific fluorescence delineation during routine surgical procedures. Future work will include efficacy demonstrations in orthotopic prostate cancer models along with comprehensive pharmacology and toxicology profiling. Additionally, integrate of the optimal 800 nm PSMA-2 prostate cancer targeted probe with a spectrally distinct, 700 nm nerve-specific fluorophore for a two-color FGS approach would benefit intraoperative decision making during RARP.^[56] [57]^ This two-color imaging strategy would offer a complete solution to improve cancer identification and adjacent critical nerve tissue visualization, simultaneously ensuring negative margins and nerve tissue preservation with surgical precision, enhancing overall patient outcomes.

## Acknowledgements

We would like to thank Laurie Akuburio, Cam Nguyen, and Nick Ferguson for experimental assistance. This work was funded by the National Cancer Institute (R21CA246413, Niedre; R01CA271532, Gibbs), a Kuni Foundation Discovery Grant (Gibbs/Wong/Niedre), and the OHSU Foundation.

