## Supplementary material for "Engineering NIR probes to enhance affinity and clinical workflow compatibility for prostate cancer imaging": SI

**Supplementary Information includes:**

**Figure S1**: *Validation of recombinant and biotinylated PSMA by SDS-PAGE and Western blot.*
**Figure S2**: *In vitro stained with PSMA-4 & a Cy7 fluorophore without the EuK ligand.*
**Materials and Methods**
**Synthetic details**
**NMR and HPLC-MS**

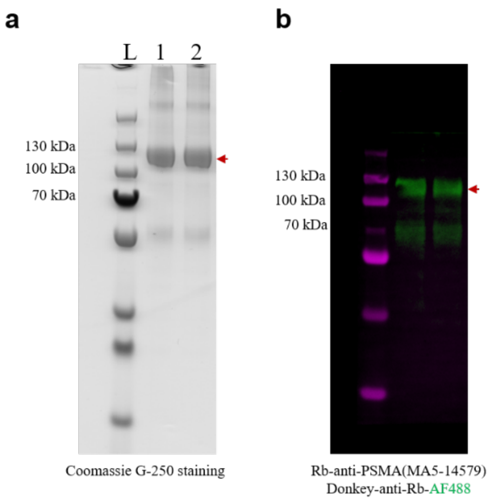


**<u>Figure S1</u>:** *Validation of recombinant and biotinylated PSMA by SDS-PAGE and Western blot.* **a**. PSMA (Expected ~84kDa) and biotinylated $^{b}$PSMA displayed in a 4-12% bis-tris gel and stained with Coomassie Brilliant Blue G-250. **b**. Fluorescent western blot confirmed the PSMA (lane 1) and $^{b}$PSMA (lane 2) bands near 100 kDa, displayed in 488 nm (green, red arrow) channel, with the protein ladder in the 647 nm (magenta) channel. The apparent molecular weight of PSMA was higher than expected, due to glycosylation modification. The blot was developed with Rabbit anti-PSMA antibody (MA5-14579). The fluorescent images above were collected using an iBright (FL1000) imaging system.

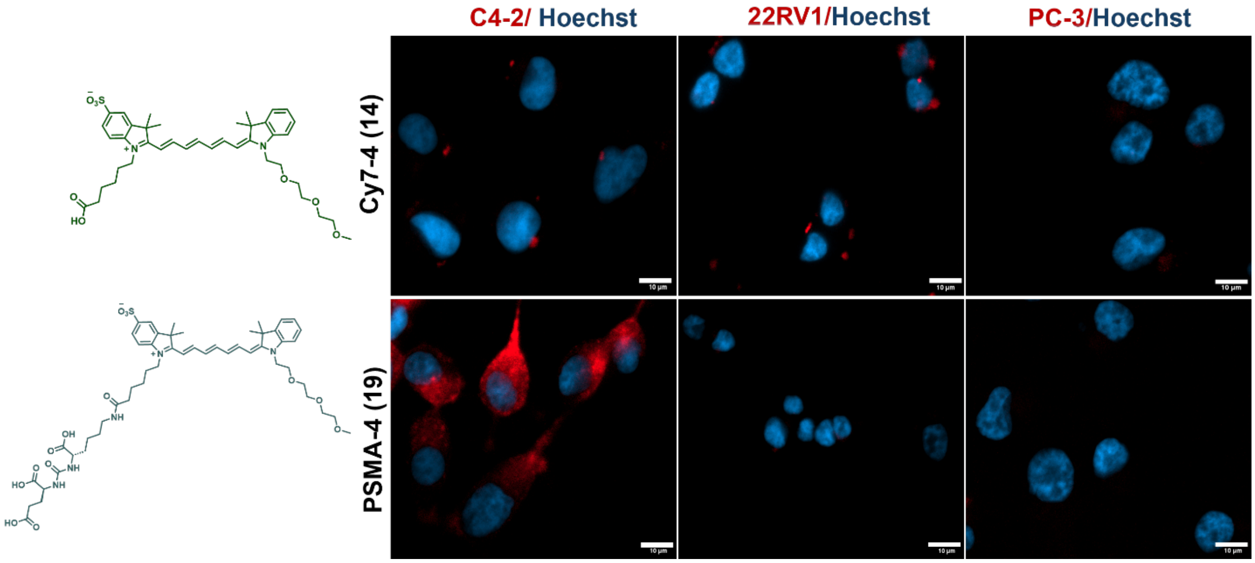


**<u>Figure S2</u>:** *In vitro stained with PSMA-4 & a Cy7 fluorophore without the EuK ligand*. Live cell images of C4-2 (PSMA++), 22RV1 (PSMA+), and PC-3 (PSMA-) prostate cancer cells stained with (**14**) Cy7-4 (top) or (**19**) PSMA-4 (bottom) at a concentration of 10 µM for 1 h at 37 °C. Cell nuclei were stained with Hoechst (blue) for 15 min. The excitation and emission filter used for imaging PSMA probes and fluorophore were (670-750)Ex/(772-855)Em. Scale bar = 10 µm.

### Materials and Methods

#### *General*

All the solvents and chemicals were ACS grade, obtained from commercially available resources and were used without further purification. HPLC-grade solvents were purchased from Fisher Scientific. Analytical TLC was performed on silica gel TLC Silica Gel 60 $F_{254}$. Purification was performed on a Biotage Selekt System using pre-packed silica gel cartridges or on a reverse phase preparative HPLC (Agilent 1250 Infinity HPLC).

#### *Prep-HPLC purification*

HPLC purification of the compounds was performed on reverse phase preparative HPLC (Agilent 1250 Infinity HPLC). The fluid handling unit was equipped with a Preparative C18 column (InfinityLab Poroshell 120 HPH-C18 Column, 21.2 x 150 mm, 4 μm), a dual-wavelength absorbance detector (254 nm, 750 nm), and a manual injector outfitted with a semi-preparative flow cell. Solvent A was water with 0.1% formic acid and solvent B was acetonitrile with 0.1% formic acid. The purification utilized a linear gradient from 10% to 90% solvent B over 50 min. Mobile phase flow rate was 10 mL/min.

#### *Structure and purity characterization*

The purity of all the compounds was measured using 1260 infinity HPLC integrated with electrospray ionization (ESI) single quadrupole (SQ) mass spectrometry (Infinity Lab LC/MSD iQ, Agilent). Analyses were carried using Agilent C18 column (InfinityLab Poroshell 120 EC-C18, 2.1 x 50 mm, 2.7 μm). The mobile phase was water with 0.1% formic acid and acetonitrile with 0.1% of formic acid with a linear gradient. The flow rate was 0.4 mL/min, and all compounds were identified by mass-to-charge (m/z) ratio. The $^{1}H$ and $^{13}C$ NMR spectra were recorded on Bruker 400 MHz Avance II+ spectrometer. MestReNova software (14.3.3-33362) was used for NMR data processing. The samples were dissolved in spectral grade $CD_3OD$-$d_4$, and DMSO-$d_6$ solvents. Chemical shifts (δ) are given in parts per million (ppm) referenced to the residual solvent peak (DMSO-$d_6$: 2.50 ppm for $^{1}H$, and 39.52 ppm for $^{13}C$; $CD_3OD$: 3.31 ppm for $^{1}H$, and 49.00 ppm for $^{13}C$). Multiplicity of each peak, singlet (s), doublet (d), triplet (t), quartet (q), doublet of doublets (dd), multiplet (m), and integration are reported for $^{1}H$ NMR.

#### *Molecular modeling and docking*

Molecular docking studies were performed using *in silico* modeling to define the binding modes of the probes with the varied PK profile modulators. X-ray structures of PSMA (PDB ID 2XEG) were retrieved from the protein data bank server. The Schrödinger Maestro interface was used for the molecular docking study.

#### *Absorption and fluorescence spectroscopy*

All probes were solubilized in phosphate buffered saline (PBS, pH 7.4) containing 1% dimethyl sulfoxide (DMSO) at a concentration of 10 μM. Absorption and fluorescence emission spectra were collected using a SpectraMax M5 spectrometer with a Microplate reader (Molecular Devices, Sunnyvale, CA). All absorption spectra were reference corrected. Extinction coefficients for all screened fluorophores were calculated using Beer's Law plots of absorbance versus concentration. Relative quantum yields were calculated using HITCI (1,10,3,3,30,30-hexamethylindotricarbocyanine iodide) as reference standard. [1] Data analyses were carried out using GraphPad Prism version 10.2.3.

***In vitro biotinylation of recombinant PSMA protein***

DNA fragment encoding the extracellular domain of PSMA (residues K44-A750, NCBI accession NM_004476) with a N-terminal AviTag and C-terminal 6xHis purification tag, was synthetically produced after codon optimization, and assembled into a pcDNA3.4 for expression in Chinese Hamster Ovary cells at GenScript (NJ, USA). Purified recombinant proteins (>80% purity) were stored in PBS (pH7.2). Avi-tagged PSMA was buffer exchanged using a 0.5 mL, 3 kDa Amicon Ultra centrifugal filter (Sigma-Aldrich) into 10 mM Tris-HCl (pH 8.0), 150 mM potassium glutamate and biotinylated with BirA biotin-protein ligase reaction kit (BirA500, Avidity) according to the manufacturer's protocol. To remove excess d-biotin 0.5 mL, 10 kDa Amicon Ultra centrifugal filters (Sigma-Aldrich) were used and buffer exchanged into DPBS (ThermoFisher). The biotinylated $^{b}$PSMA was further validated by the SDS-PAGE and western blot analysis according to previously established protocols (Fig. S1). [2] All samples were prepared by mixing NuPAGE® LDS sample buffer (4X) (ThermoFisher) with 50 mM DTT (Sigma-Aldrich), followed by heating the sample for denaturing electrophoresis at 95 $^{o}$C for 10 min. SDS-PAGE was performed using NuPAGE® 4-12% Bis-Tris mini protein gel (ThermoFisher) in NuPAGE® MES SDS running buffer (ThermoFisher). For direct gel staining, SimpleBlue® safestain (ThermoFisher) was used according to manufacturer's protocol. For western blot analysis, protein samples were transferred to a 0.45 µm Immobilon®-P PVDF membrane in NuPAGE® transfer buffer. After washing with DPBS, the membrane was blocked for 1 h at room temperature using intercept® PBS (LI-COR Biosciences). The membrane was then incubated with rabbit anti-PSMA antibody (ThermoFisher) in blocking buffer with 0.2% Tween-20 for 1 h at room temperature. Following three washes with DPBS, the membrane was further developed with donkey anti-rabbit-AF488 (ThermoFisher) in blocking buffer with 0.2% Tween-20 for 1 h at room temperature. Following three washes with DPBS, membranes were visualized on an iBright imaging system (ThermoFisher).

***Bio-layer Interferometry (BLI) for affinity determination***

Kinetic assays were performed using a ForteBio Octet® RED384 instrument at 30 $^{o}$C with data acquisition software 9.0, following established protocols. [2] Assays were performed in 384-well tilted-bottom plates (Sartorius) with orbital shaking at 1,000 rpm. The running buffer contained DPBS (pH 7.4), 0.1% bovine serum albumin (BSA), 0.1% DMSO and 0.01% Tween-20. Streptavidin (SA) biosensors (Sartorius) pre-equilibrated in DPBS for 10 min, then loaded with 50 µg/mL of biotinylated PSMA ligand or biocytin (Sigma-Aldrich) as control for 20 min. Sensors were blocked with 100 μg/mL of biocytin for 5 min, washed in running buffer for 1 min, and equilibrated in running buffer for 5 min to establish baseline. Binding of PSMA probe analytes (0-0.8 μM), were assessed with a 90 s association and a 60 s dissociation steps. Biocytin-loaded sensors served as parallel references to correct for background. Data were processed using ForteBio Data Analysis HT 10.0 by subtracting reference sensor (biocytin loaded) and reference well (no analyte) signals. Binding kinetics were globally fit to a 1:1 Langmuir model using at least four different concentrations, across three independent experiments. Association ($k_a$) and dissociation ($k_d$) rate constants were averaged, and the equilibrium dissociation constant ($K_D$) was calculated as $k_d / k_a$. Final data were exported and plotted in GraphPad Prism9.

***General cell culture***

The human prostate cancer cell lines, C4-2 (ATCC, CRL 3314), 22RV1 (ATCC, CRL 2505) and PC-3 (ATCC, CRL-1435) were purchased from ATCC. The cell lines were expanded in their respective optimal growth media C4-2, PC-3: DMEM (Gibco) + 10% fetal bovine serum (FBS, Cytiva) + 1% Penicillin-Streptomycin-glutamine (Gibco); 22RV1: RPMI 1640 (Gibco) + 10% fetal bovine serum + 1% Penicillin-Streptomycin-glutamine and stored at 37 °C in a 5% $CO_2$ incubator. All cells were maintained mycoplasma-free at passage numbers <25 for all studies. PSMA antibody (Mouse Hybridoma Prost J533) was purchased from ATCC, HB-12127.

***Live-cell imaging***

For live-cell fluorescence microscopy experiments, C4-2 cells, PC-3 cells were passaged in DMEM with FBS (10%) + 1% Penicillin-Streptomycin-glutamine and 22RV1 cells were passaged in RPMI with FBS (10%) + 1% Penicillin-Streptomycin-glutamine and plated in 8-well plates with 1.5H cover glass bottoms. 24 h later, the media was removed, and cells were incubated in DMEM (FluoroBrite, Gibco) containing 10 μM of PSMA-1, PSMA-2, PSMA-3, or PSMA-4 at 37 °C for 1 h. PSMA antibody J533 was added at a concentration of 3 nM and incubated at 37 °C for 1 h. Staining media was removed and fresh media containing Hoechst 33342 was added. Cells were incubated for an additional 10 min, then washed with FluororBrtite DMEM media for 3 × 5 min. Cells were then washed with PBS 1 × 5 min. Cells were incubated with 4% paraformaldehyde (PFA, Electron Microscopy Sciences) in PBS for another 15 min at room temperature, then washed with PBS (3 × 5 min). Fixed cells were stained with Alexa Fluor 546 phalloidin (ThermoFisher Scientific) and incubated for 20 min at 4 °C, then washed with PBS 3 × 5 min. Cells were incubated with 4% PFA in PBS for another 15 min at room temperature, then washed with PBS (3 × 5 min) before fluorescence images were acquired on an Apotome-3 (Zeiss). The following excitation and emission filter combinations were utilized for image collection: AF488 (450-490)Ex/(500-550)Em, AF546 (538-562)Ex/(570-640)Em, and AF750 (670-750)Ex/(772-855)Em. Images were processed using ImageJ/FIJI.

***Single-cell fluorescence intensity quantification***

To generate the scatter plot, images from each cell line were first loaded and processed. For each image, cells were segmented using the Mesmer algorithm, where the Hoescht channel was used as the nuclear input and the actin stain channel as the membrane input to predict cell boundaries. This yielded labeled masks for the nucleus and the whole cell. A cytoplasm mask was then created by subtracting the nuclear mask from the whole cell mask on a pixel-by-pixel basis. For each cell identified in the cytoplasm mask, the average intensity of the test probe channel was calculated and plotted as an individual point on the scatter plot. Additionally, the average intensity and standard deviation across all cells were computed, and the mean value was displayed with error bars representing ±1 standard deviation. The final plot presents data for three different cell lines, each showing a cluster of points corresponding to per-cell average intensities and an overlay of the mean ± standard deviation.

***Blood PK assessment by Diffuse in vivo flow cytometry (DiFC)***

Diffuse *in vivo* Flow Cytometry (DiFC) was performed to noninvasively assess blood pharmacokinetic (PK) profiles of the four PSMA targeted probes. [3] The instrument contained a 730 nm laser (0730-06-01-0050-100; Cobolt) with a 720/24nm excitation filter (BP1-ex; FF01-720/24-12.5, IDEX Health and Science LLC) for Cy7 excitation with the light power at the sample tuned to 25 mW using a neutral density (ND) filter. The laser was coupled into a source fiber with a collimation package (FC-ex; F240SMA-633, Thorlabs Inc., Newton, NJ). The output of the collection fibers was collimated (FC-em; F240SMA-633, Thorlabs) and the light was passed through an 780 nm longpass emission filter for Cy7 detection (LP-em; ET780lp, Chroma, Bellows Falls, VT) before being focused onto the surface of a photomultiplier tube (PMT; H10722-20, Hamamatsu, Bridgewater, New Jersey) with a 30 mm focal length lens (L-em; 45363, Edmund Optics). The PMT was powered by a power supply (C10709, Hamamatsu). Output signals from the PMTs were filtered with an electronic 100 Hz low pass filter, amplified with a low-noise voltage pre-amplifier (PA; SR560, Stanford Research Systems, Sunnyvale, CA), and then acquired with a data acquisition board (DAQ; USB-6343 BNC; National Instruments, Austin, TX). DiFC uses a custom-designed integrated optical fiber sensor assembly (EmVision LLC, Loxahatchee, FL). The optical fiber sensor is constructed with 7 all silica low hydroxyl (OH) content 300 $\mu$m core 0.22 NA collection fibers, for Cy7 detection (780-810 nm collection filter). The 7 collection fibers are arranged around a 730 nm laser delivery fiber with a 697/75nm excitation filter which is also an all silica 300 $\mu$m more low OH, 0.22 NA fiber.

***Animal care and use***

All DiFC mice and experiments were handled in accordance with Northeastern University's Institutional Animal Care and Use committee (IACUC) policies on animal care, protocol #24-0207R. 6-8-week-old male Athymic nude mice (Athymic NCR Nu/Nu/553Charles River Labs, Cambridge, MA) were used for this study. Mice were fed a diet of low fluorescence animal chow (AIN 93M Mature Rodent Diet, Ziegler Feed, East Berlin, PA). Mice were maintained under inhaled isoflurane during Red-NIR-DiFC scanning to prevent movement and were kept warm with a heating pad under the body. All *in vivo* fluorescence imaging animal experiments were conducted with the approval of the Institutional Animal Care and Use Committee (IACUC) at Oregon Health and Science University (OHSU). All mice were housed and handled following an IACUC-approved protocol. Male mice were used to assess tumor imaging with PSMA-1, PSMA-2, PSMA-3, and PSMA-4. At the end of the imaging study, the mice were humanely euthanized with carbon dioxide inhalation as the primary method, followed by cervical dislocation as a confirmatory procedure.

***Imaging and quantification (DiFC experiment)***

The saphenous vessels in the hindlimb were scanned by placing the optical fiber probe on the skin over the vessels with ultrasound gel applied for improved light delivery and collection. The instrument PMT gain was adjusted such that the signal coming from an uninjected mouse was approximately 1-2 mV (baseline). After the initial adjustment, the gain was kept at the same value for all experiments. Five minutes of baseline fluorescence data was collected for each mouse, after which 10 nmol of PSMA-1, PSMA-2, PSMA-3, or PSMA-4 was administered via retroorbital (R.O.) injection (n=4 mice/probe). Mice were then scanned until the measured fluorescence signal returned to the baseline value as measured prior to injection. The collected fluorescence data for each mouse was averaged every 5 min and normalized. Using the averaged and normalized data for each mouse, the average and standard deviation was calculated for the 4 mice used to test each PSMA probe.

***Tumor Implantation and Mouse Xenograft models***

The human prostate cancer cells, C4-2 were grown to 95% confluence for xenograft implantation. C4-2 cells were implanted in Matrigel (Corning) with cold DMEM in 1:1 ratio at a concentration of 3 X $10^6$ cells/100 µL. Athymic nude mice were anesthetized with 2% isoflurane, and the cells were implanted bilaterally into the flank region using a 25-gauge needle. Tumor growth was monitored and restricted to a maximum size of 0.75 $cm^3$ in accordance with the approved IACUC protocol. Imaging studies were conducted on tumors measuring between 0.5 and 0.75 $cm^3$.

***Imaging and Quantification***

All images were acquired using a custom-built small animal imaging system.[4] [4b, 5] [6] [7] [7b] This system enabled real-time color and fluorescence imaging and featured a QImaging EXi Blue monochrome camera (Surrey, British Columbia, Canada) for fluorescence detection. A removable Bayer filter allowed for the collection of co-registered color and fluorescence images. White light illumination was provided by a PhotoFluor II light source (89 North, Burlington, Vermont, USA), which was directed onto the surgical field through a liquid light guide and used unfiltered. For fluorescence imaging of PSMA-targeted probes, the PhotoFluor II light source was filtered with a 740 ± 20 nm bandpass excitation filter, while fluorescence emission was captured using a 780 nm long-pass filter. To ensure quantitative comparison, camera and light source positions remained fixed throughout the imaging studies. Camera exposure times varied from 2-800 ms, with all images normalized for exposure time. Tissue fluorescence intensities were quantified using custom-written MATLAB code, which enabled the selection of regions of interest (ROIs) for muscle and tumor tissues on white light images while remaining blinded to fluorescence images. These selected ROIs were then superimposed onto the corresponding fluorescence images, allowing for unbiased quantification of tissue fluorescence intensities and the calculation of tumor-to-muscle (T/M) and signal-to-background ratios (SBRs).

### Synthetic Details

***Structures of intermediates (compounds 7-10)***

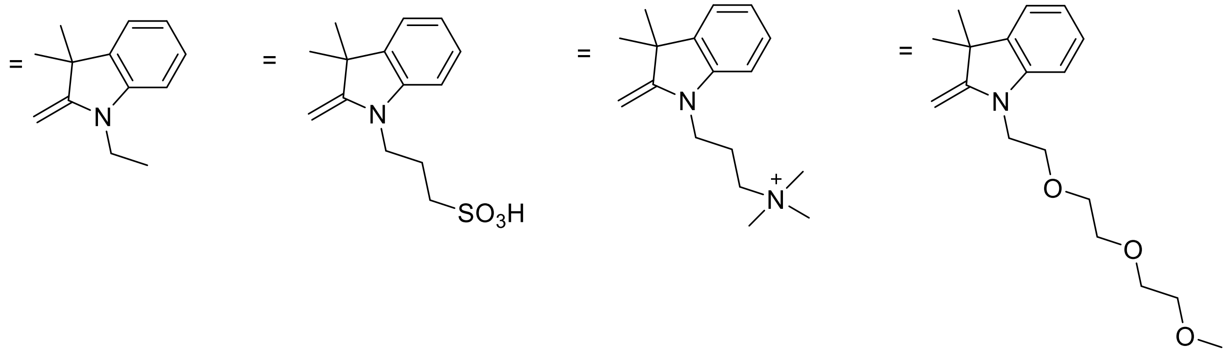

***Synthesis of compounds 11-14***

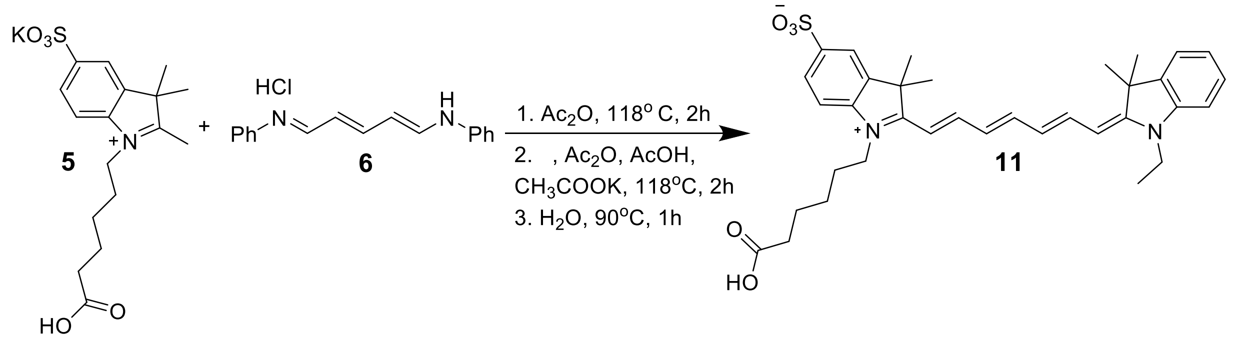

**Scheme S2.** Synthetic route to intermediate **11**.

**1-(5-carboxypentyl)-2-((1E,3E,5E)-7-((E)-1-ethyl-3,3-dimethylindolin-2-ylidene)hepta-1,3,5-trien-1-yl)-3,3-dimethyl-3H-indol-1-ium-5-sulfonate, 11:** Compound **5** (0.250 g, 0.636 mmol, 1 equiv), N-[5-(Phenylamino)-2,4-pentadienylidene]aniline monohydrochloride **6** (0.181 g, 0.636 mmol, 1 equiv), acetic anhydride (6 mL) were combined and refluxed at 118 °C for 2 h. Compound **7** (0.120 g, 0.636 mmol, 1 equiv), potassium acetate (0.312 g, 3.18 mmol, 5 equiv), acetic anhydride (2.5 mL) and acetic acid (1.5 mL) were added to the reaction mixture and stirred 118 °C for an additional 2 h. The reaction temperature was then reduced to 90 °C and 2.5 mL of water was added and stirred for another 1 h. The reaction mixture was then cooled to room temperature, the solvent was removed under reduced pressure. Compound **11** was purified by prep HPLC using a gradient of MeCN-$H_2O$ with 0.1% of formic acid. The purified product was obtained as a green solid (0.074 g, 20% yield). $^1$H NMR (400 MHz, DMSO-$d_6$) δ 12.02 (s, 1H), 7.91-7.86 (m, 2H), 7.80 (d,1H) 7.73 (s, 1H), 7.61(dd, 2H), 7.40 (d, 2H), 7.26 (t, 2H), 6.56 (dd, 2H), 6.43 (d, 1H), 6.33 (d, 1H), 4.15 (d, 2H), 4.04 (t, 2H), 2.20 (t, 2H), 1.68 (d, 2H), 1.63 (d, 12H), 1.54 (t, 2H), 1.39-1.35 (m, 2H), 1.27 (t, 3H). $^{13}$C NMR (101 MHz, DMSO-$d_6$) δ 174.30. 171.99, 171.53, 170.65, 156.27, 151.78, 150.57, 144.76, 142.28, 141.60, 141.26, 140.11, 128.52, 126.11, 125.48, 124.86, 122.49, 119.74, 111.02, 109.80, 104.05, 103.48, 54.94, 48.87, 48.40, 43.34, 33.52, 27.20, 27.03,

26.67,25.65, 24.24, 22.94, 21.09, 12.31. LC-MS(ESI) m/z $[M+H]^+$ found 603.2, calcd for $C_{35}H_{43}N_2O_5S^+$ 603.28.

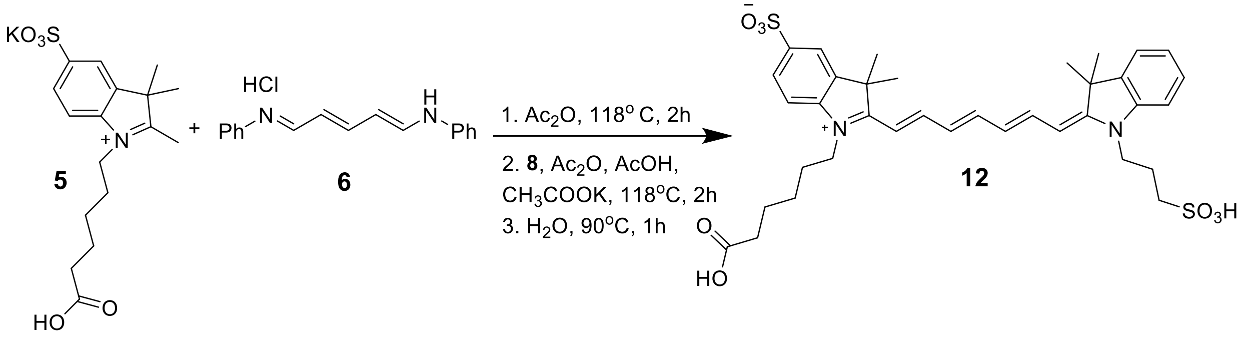


**Scheme S3.** Synthetic route to intermediate **12**.

**1-(5-carboxypentyl)-2-((1*E*,3*E*,5*E*)-7-((*E*)-3,3-dimethyl-1-(3-sulfonatopropyl)indolin-2-ylidene)hepta-1,3,5-trien-1-yl)-3,3-dimethyl-3*H*-indol-1-ium-5-sulfonate, 12**: Compound **5** (0.500 g, 1.273 mmol, 1 equiv), N-[5-(Phenylamino)-2,4-pentadienylidene]aniline monohydrochloride **6** (0.363 g, 1.27 mmol, 1 equiv), acetic anhydride (12.7 mL) were combined and refluxed at 118 °C for 2 h. Compound **8** (0.357 g, 1.27 mmol, 1 equiv), potassium acetate (0.623 g, 6.35 mmol, 5 equiv), acetic anhydride (5 mL) and acetic acid (2.5 mL) were added to the reaction mixture and stirred at 118 °C for an additional 2 h. The reaction temperature was then reduced to 90 °C and 5 mL water was added stirred for 1 h. The reaction mixture was then cooled to room temperature, the solvent was removed under reduced pressure. Compound **12** was purified by prep HPLC using gradient of MeCN-$H_2O$ with 0.1% of formic acid. The purified product was obtained as a green solid (0.071 g, 8% yield). $^1H$ NMR (400 MHz, DMSO- $d_6$) δ 12.06 (s, 1H), 7.95-7.78 (m, 4H), 7.71 (S, 1H),7.63-7.52 (M, 2H), 7.51 (d, 1H) 7.40 (t, 1H), 7.25 (t, 2H), 6.57-6.48 (m, 4H), 6.31 (d, 1H), 4.27 (t, 2H), 4.02 (t, 2H), 2.58 (t, 2H), 2.13-2.08 (m, 3H), 2.02-1.99 (m, 3H), 1.71-1.05 (m, 2H), 1.63 (s, 12H), 1.57-1.49(m, 3H), 1.40-1.23(m, 4H). $^{13}C$ NMR (101 MHz, DMSO- $d_6$) δ 174.37, 172.15, 170.50, 156.25, 152.02, 150.49, 144.49, 142.37, 142.09, 141.23, 140.13, 130.24, 128.54, 128.16, 126.12, 124.76, 122.43, 119.73, 111.38, 109.75, 104.61,103.38, 48.93, 48.39, 47.94, 43.31, 42.79, 33.53, 27.25, 27.12, 25.67, 26.66, 24.25, 23.54. LC-MS(ESI) m/z $[M-H]^-$ found 695.2 calcd for $C_{36}H_{43}N_2O_8S_2^-$ 695.24.

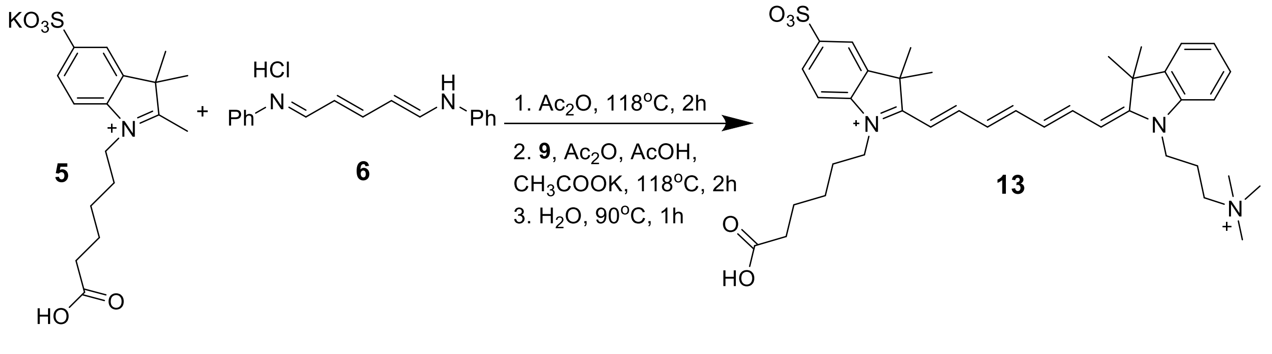

**Scheme S4.** Synthetic route to intermediate **13**.

**1-(5-carboxypentyl)-2-((1*E*,3*E*,5*E*)-7-((*E*)-3,3-dimethyl-1-(3-(trimethylammonio)propyl)indolin-2-ylidene) hepta-1,3,5-trien-1-yl)-3,3-dimethyl-3*H*-indol-1-ium-5-sulfonate, 13:** Compound **5** (0.100 g, 0.254 mmol, 1 equiv), N-[5-(Phenylamino)-2,4-pentadienylidene]aniline monohydrochloride **6** (0.072 g, 0.254 mmol, 1 equiv), acetic anhydride (2.5 mL) were combined and refluxed at 118 °C for 2 h. Compound **9** (0.066 g, 0.254 mmol, 1 equiv), potassium acetate (0.125 g, 1.27 mmol, 5 equiv), acetic anhydride (1 mL) and acetic acid (0.5 mL) were added to the reaction mixture and stirred at 118 °C for an additional 2 h. The reaction temperature was then reduced to 90 °C and 1 mL of water was added and stirred for another 1 h. The reaction mixture was then cooled to room temperature, the solvent was removed under reduced pressure. Compound **13** was purified by prep HPLC using a gradient of MeCN-$H_2O$ with 0.1% of formic acid. The purified product was obtained as a green solid (0.018 g, 11% yield). $^1$H NMR (400 MHz, DMSO- *d*$_6$) δ 11.77 (s, 1H), 7.97-7.85 (m, 2H), 7.78-7.74 (dd, 2H), 7.64 (t, 1H), 7.55 (d, 1H), 7.39-7.36 (m, 2H), 7.31-7.29 (m, 1H), 4.12-4.04 (m, 4H), 3.52 (s, 2H), 3.07 (s, 9H), 2.20-2.17 (m, 2H), 2.12-2.09 (m, 2H), 1.64 (s, 12H), 1.56-1.52 (m, 2H), 1.41-1.35 (m, 4H). $^{13}$C NMR (101 MHz, DMSO-*d*$_6$) δ 174.81, 171.93, 170.47, 146.10, 145.60, 142.62, 142.55, 142.42, 141.14, 140.79, 132.53, 130.15, 128.88, 126.60, 126.20, 124.68, 122.89, 120.27, 110.94, 110.56, 62.89, 52.92, 49.47, 49.12, 34.00, 27.81, 27.61, 27.47, 26.12, 24.71, 23.41, 21.17. LC-MS(ESI) m/z $[M+H]^+$ found 674.7, $[M+2H]^+$ 337.9, calcd. for $C_{39}H_{53}N_3O_5S^{2+}$ 337.68.

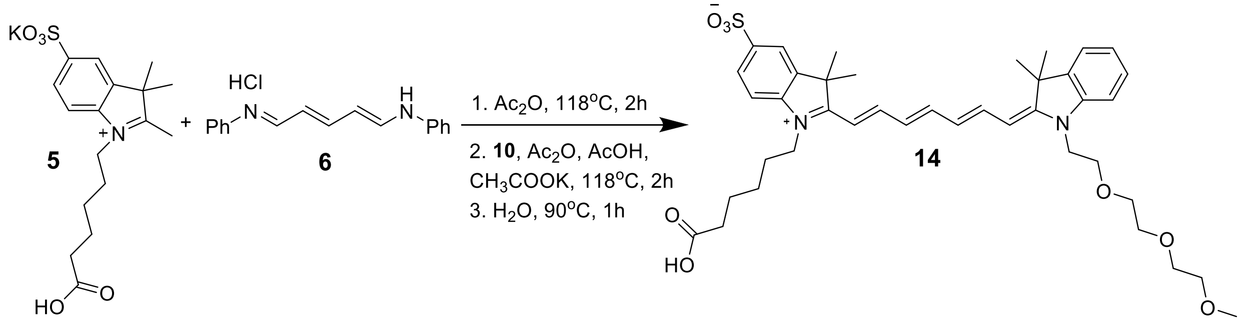


**Scheme S5.** Synthetic route to intermediate **14**.

**1-(5-carboxypentyl)-2-((1*E*,3*E*,5*E*)-7-((*E*)-1-(2-(2-(2-methoxyethoxy)ethoxy)ethyl)-3,3-dimethylindolin-2-ylidene)hepta-1,3,5-trien-1-yl)-3,3-dimethyl-3*H*-indol-1-ium-5-sulfonate, 14:** Compound **5** (0.500 g, 1.273 mmol, 1 equiv), N-[5-(Phenylamino)-2,4-pentadienylidene]aniline monohydrochloride **6** (0.363 g, 1.27 mmol, 1 equiv), acetic anhydride (12.7 mL) were combined and refluxed at 118 °C for 2 h. Compound **10** (0.388 g, 1.27 mmol, 1 equiv), potassium acetate (0.624 g,6.37 mmol, 5 equiv), acetic anhydride (5 mL) and acetic acid (2.5 mL) were added to the reaction mixture and stirred at 118 °C for an additional 2 h. The reaction temperature was then reduced to 90 °C and 5 mL of water was added and stirred for another 1 h. The reaction mixture was then cooled to room temperature, the solvent was removed under reduced pressure. Compound **14** was purified by prep HPLC using a gradient of MeCN-$H_2O$ with 0.1% of formic acid. The purified product was obtained as a green solid (0.090 g, 10% yield). $^1H$ NMR (400 MHz, $CD_3OD$-$d_4$) δ 8.04 (t, 1H), 7.84-7.82(m, 3H), 7.60 (t, 1H), 7.53 (d, 1H), 7.42 (m, 2H) 7.31 (m,1H), 6.61-6.51 (m, 3H), 6.16 (d, 1H), 4.40 (t, 2H), 3.98 (t, 2H), 3.90 (t, 2H), 3.59-3.56 (m,2H), 3.51-3.50 (m-2H), 3.48-3.46(m,2H), 3.41-3.38(m, 2H), 3.31-3.30(m, 2H) 3.29 (s,3H), 2.30 (t, 2H), 1.80-1.76 (m, 2H), 1.72 (s, 6H), 1.68()s,6H), 1.50-1.45(m, 2H). $^{13}C$ NMR (101 MHz, $CD_3OD$- $d_4$) δ 175.11, 174. 59, 168.57, 155.42, 152.67, 143.42, 141.57, 140.77, 139.63, 127.60, 125.84, 124.86, 121.24, 119.09, 111.18, 108.25, 70.80, 69.92, 69.51, 69.31, 67.03, 56.98, 48.96, 46.94, 46.87, 46.73, 46.66, 46.45, 46.23, 43.84, 32.52, 26.04, 25.75, 25.72, 25.21, 23.60. LC-MS(ESI) m/z $[M+H]^+$ found 721.3, calcd. for $C_{40}H_{53}N_2O_8S^+$ 721.35.

***Synthesis of the PSMA probes (compounds 16-19)***

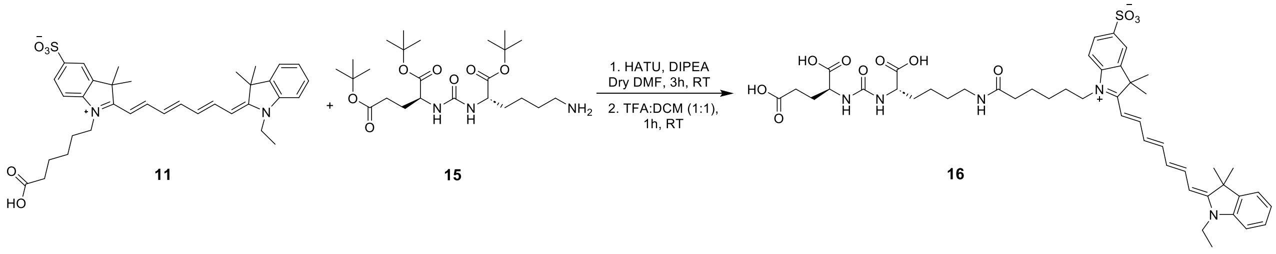


**Scheme S6.** Synthetic route to compound **16**.

**(((*S*)-1-carboxylato-5-(6-(2-((1*E*,3*E*,5*E*)-7-((*E*)-1-ethyl-3,3-dimethylindolin-2-ylidene)hepta-1,3,5-trien-1-yl)-3,3-dimethyl-5-sulfonato-3*H*-indol-1-ium-1-yl)hexanamido)pentyl)carbamoyl)-*L*-glutamate, 16:** Compound **11** (0.026 g, 0.0437 mmol, 1 equiv) and HATU (0.0183 g, 0.0481 mmol, 1.1 equiv) were dissolved in anhydrous DMF (1.2 mL). The reaction mixture was stirred for 5 min. DiPEA (19 µL, 0.109 mmol, 2.5 equiv) was added to the reaction mixture and stirred for an additional 10 min. Compound **15** (0.0316 g, 0.065 mmol, 1.5 equiv) was then added to the reaction mixture and stirred for 3 h. The reaction mixture was then acidified by adding an excess of formic acid. The reaction mixture was concentrated under reduced pressure and passed through a C18 cartridge. The product was subsequently eluted using acetonitrile and methanol. The solvent was removed under reduced pressure, and the resulting crude product was dissolved in DCM and TFA mix (1:1, 1 mL) and stirred for 1 h at room temperature. After confirming reaction completion, the solvent was removed under reduced pressure and compound **16** was purified by prep-HPLC using a gradient of MeCN-$H_2O$ with 0.1% of formic acid. The purified product was obtained as a green solid (0.010 g, 25% yield). $^1H$ NMR (400 MHz, DMSO-$d_6$) δ 12.61 (s, 3H), 7.93-7.76 (m, 2H), 7.74-7.72 (m, 2H), 7.64-7.58 (m, 2H) 7.41-7.38 (m, 2H), 7.28-7.22 (m, 2H), 6.54 (t, 2H), 6.42-6.31 (m, 4H), 4.16-4.08 (m, 2H), 4.07-3.99 (m, 3H), 2.99-2.94 (m, 2H), 2.24 (t, 2H), 2.03 (t, 2H), 1.88-1.83 (m, 1H), 1.76-1.71 (m, 2H), 1.63 (s, 12H), 1.55-1.52 (m, 3H), 1.33-1.22 (m, 6H). $^{13}C$ NMR (101 MHz, DMSO- $d_6$) δ 175.07, 174.69, 174.32, 172.14, 157.76, 145.34, 142.71, 142.12, 141.71, 140.62, 128.98, 126.61, 125.26,122.96, 120.23, 111.41, 110.37, 52.79, 52.31, 49.30, 48.95, 30.83, 28.31, 29.31, 27.66,

27.18, 27.553, 26.23, 25.41, 27.53, 26.23, 25.41, 23.09, 12.75. LC-MS m/z $[M+H]^+$ m/z found 904.2, $[M+2H]^+$ m/z 453, calcd. for $C_{47}H_{62}N_5O_{11}S^+$ 904.41.

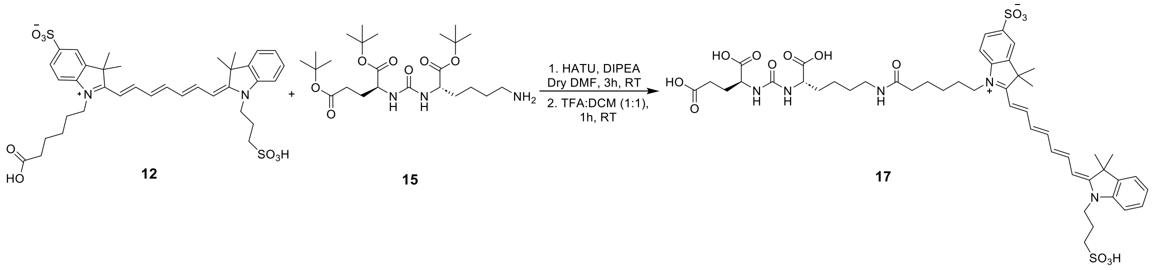


**Scheme S7.** Synthetic route to compound **17**.

**(((*S*)-1-carboxylato-5-(6-(2-((1*E*,3*E*,5*E*)-7-((*E*)-3,3-dimethyl-1-(3-sulfonatopropyl)indolin-2-ylidene)hepta-1,3,5-trien-1-yl)-3,3-dimethyl-5-sulfonato-3*H*-indol-1-ium-1-yl)hexanamido)pentyl)carbamoyl)-*L*-glutamate, 17:** Compound **12** (0.031 g, 0.044 mmol, 1 equiv) and HATU (0.0180 g, 0.048 mmol, 1.1 equiv) were dissolved in anhydrous DMF (1 mL). The reaction mixture was stirred for 5 min. DiPEA (19 µL, 0.11 mmol, 2.5 equiv) was added to the reaction mixture and stirred for an additional 10 min. Compound **15**, (0.032 g, 0.066 mmol, 1.5 equiv) was then added to the reaction mixture and stirred for 3 h. The reaction mixture was acidified by adding an excess of formic acid. The reaction mixture was concentrated under reduced pressure and passed through a C18 cartridge. The product was subsequently eluted using acetonitrile and methanol. The solvent was removed under reduced pressure, and the resulting crude product was dissolved in DCM and TFA mix (1:1, 1 mL) and stirred for 1 h at room temperature. After confirming reaction completion, the solvent was removed under reduced pressure and compound **17** was purified by prep HPLC using a gradient of MeCN-$H_2O$ with 0.1% of formic acid. The purified product was obtained as a green solid (0.009 g, 23% yield). $^1H$ NMR (400 MHz, DMSO- $d_6$) δ 7.95-7.81 (m, 2H), 7.75-7.72 (m, 3H), 7.63-7.58 (m, 2H), 7.50(d, 1H), 7.42-7.38 (m, 1H), 7.27-7.22 (m, 2H), 6.57-6.48 (m, 3H), 6.36-6.28 (m, 3H), 4.25 (t, 2H), 4.09-4.00 (m, 6H), 2.99-2.94 (m, 2H), 2.58 (t, 2H), 2.28-2.21 (m, 2H), 2.05-1.98 (m, 3H), 1.94-1.70 (m, 2H), 1.64 (s, 12H), 1.55-1.51 (m, 4H), 1.35-1.22 (m, 6H). $^{13}C$ NMR (101 MHz, DMSO- $d_6$) δ 175.02, 174.67, 174.20, 172.18, 157.82, 144.98, 142.85, 141.68, 142.55, 140.57, 128.98, 126.59, 125.93, 122.87, 120.19, 111.83, 110.22, 52.76, 52.16, 49.37, 28.84, 48.37, 43.23, 40.89, 38.70, 35.60, 32.15, 30.39, 29.29, 27.91, 27.70, 27.57, 27.14, 26.22, 25.40, 23.99, 23.08. LC-MS(ESI) $[M+2H]^+$ m/z found 998.4, calcd for $C_{48}H_{64}N_5O_{14}S_2^+$ 998.38.

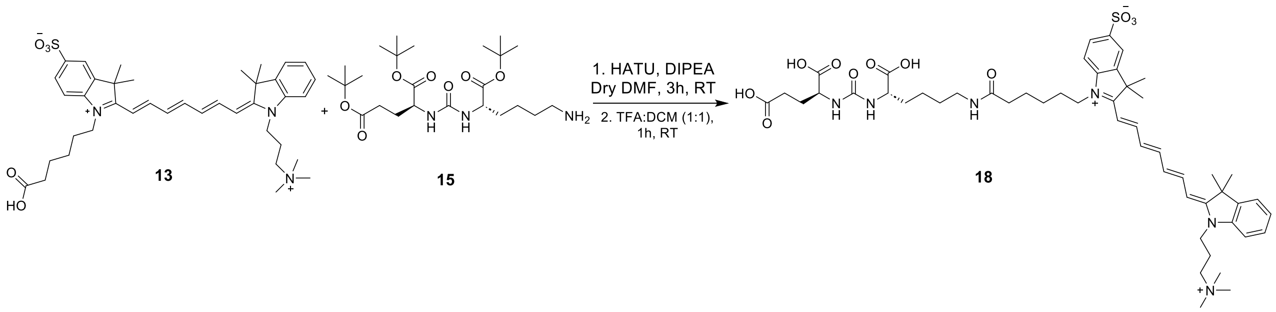

**Scheme S8.** Synthetic route to compound **18**.

**((($S$)-1-carboxylato-5-(6-(2-((1$E$,3$E$,5$E$)-7-(($E$)-3,3-dimethyl-1-(3-(trimethylammonio)propyl)indolin-2-ylidene)hepta-1,3,5-trien-1-yl)-3,3-dimethyl-5-sulfonato-3$H$-indol-1-ium-1-yl)hexanamido)pentyl)carbamoyl)-$L$-glutamate, 18:** Compound **13** (0.050 g, 0.072 mmol, 1 equiv) and HATU (0.030 g, 0.079 mmol, 1.1 equiv) were dissolved in anhydrous DMF (1.2 mL). The reaction mixture was stirred for 5 min. DiPEA (30 µL, 0.18 mmol, 2.5 equiv) was added to the reaction mixture and stirred for an additional 10 min. Compound **15**, (0.053 g, 0.108 mmol, 1.5 equiv) was then added to the reaction mixture and stirred for 3 h. The reaction mixture was then acidified by adding an excess of formic acid. The reaction mixture was concentrated under reduced pressure and passed through a C18 cartridge. The product was subsequently eluted using acetonitrile and methanol. The solvent was removed under reduced pressure, and the resulting crude product was dissolved in DCM and TFA mix (1:1, 1 mL) and stirred for 1 h at room temperature. After confirming reaction completion, the solvent was removed under reduced pressure and compound **18** was purified by prep-HPLC using a gradient of MeCN-$H_2O$ with 0.1% of formic acid. The purified product was obtained as a green solid (0.012 g, 18% yield).$^1$H NMR (400 MHz, DMSO-$d_6$) δ 7.97-7.84 (m, 2H), 7.81-7.74 (m, 3H), 7.65 (d, 1H), 7.57 (d, 1H), 7.37 (t, 3H), 6.63-6.53 (m, 2H), 6.50-6.42 (m, 3H), 6.29 (s, 1H), 4.12-3.96 (m, 6H), 3.56-3.51 (m, 2H), 3.09 (s, 9H), 3.00-2.98 (m, 2H), 2.33-2.26 (m, 2H), 2.14-2.11 (m, 3H), 2.06 (t, 2H), 1.87-1.82 (m, 2H), 1.64 (s, 12H), 1.554-1.50 (m, 4H), 1.36-1.27 (m, 6H). $^{13}$C NMR (101 MHz, DMSO-$d_6$) δ 175.39, 174.92, 174.84, 173.17, 172.20, 170.50, 164.42, 157.76, 151.09, 145.94, 142.54, 142.39, 141.18, 141.10, 128.86, 124.67, 122.90, 120.27, 111.08, 110.89, 62.92, 53.30, 52.95, 49.45, 48.83, 40.64, 38.77, 35.56, 32.49, 29.37, 29.48, 27.81, 27.26, 27.49, 26.07, 25.30, 23.17, 21.18. LC-MS(ESI) $[M+2H]^+$ 488.5, calcd for $C_{52}H_{72}N_6O_{11}S^{2+}$ 488.24.

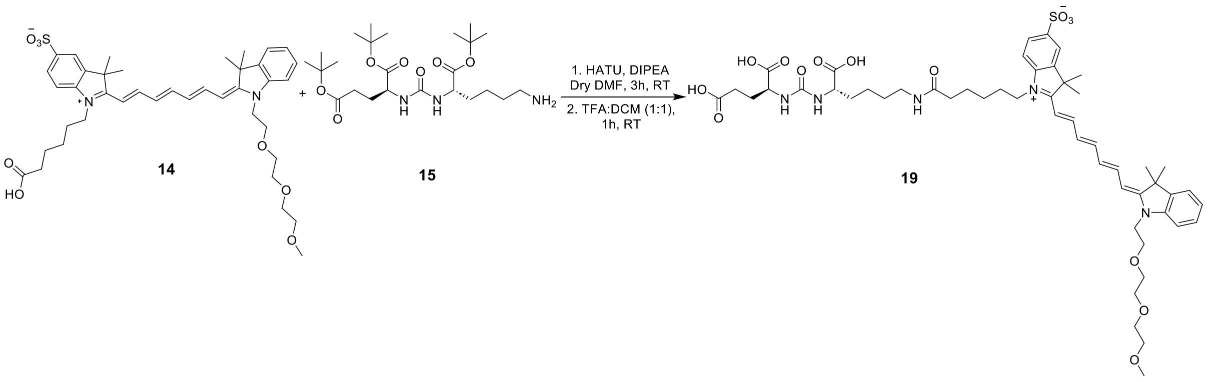


**Scheme S9.** Synthetic route to compound **19**.

**(((*S*)-1-carboxylato-5-(6-(2-((1*E*,3*E*,5*E*)-7-((*E*)-1-(2-(2-(2-methoxyethoxy)ethoxy)ethyl)-3,3-dimethylindolin-2-ylidene)hepta-1,3,5-trien-1-yl)-3,3-dimethyl-5-sulfonato-3*H*-indol-1-ium-1-yl)hexanamido)pentyl)carbamoyl)-*L*-glutamate, 19:** Compound **14** (0.079 g, 0.110 mmol, 1 equiv) and HATU (0.0460 g, 0.121 mmol, 1.1 equiv) were dissolved in anhydrous DMF (1.2 mL). The reaction mixture was stirred for 5 min. DiPEA (48.51 µL, 0.275 mmol, 2.5 equiv) was added to the reaction mixture and stirred for an additional 10 min. Compound **15** (0.080 g, 0.165 mmol, 1.5 equiv) was then added to the reaction mixture and stirred for 3 h. The reaction mixture was acidified by adding an excess of formic acid. The reaction mixture was concentrated under reduced pressure and passed through a C18 cartridge. The product was subsequently eluted using acetonitrile and methanol. The solvent was removed under reduced pressure, and the resulting crude product was dissolved in DCM and TFA mix (1:1, 1 mL) and stirred for 1 h at room temperature. The reaction mixture was then acidified by adding an excess of formic acid. After confirming reaction completion, the solvent was removed under reduced pressure and compound **19** was purified by prep HPLC using a gradient of MeCN-$H_2O$ with 0.1% of formic acid. The purified product was obtained as a green solid (0.020 g, 18% yield). $^1$H NMR (400 MHz, DMSO-*d₆*) δ 12.66 (s, 3H), 7.92-7.83 (m, 2H), 7.78-7.71 (m, 3H), 7.64 (dd, 1H), 7.58 (d, 1H), 6.58-6.48 (m, 2H), 6.44 (d, 1H), 6.37-6.32 (m, 3H), 4.30 (t, 2H), 4.10-3.99 (m, 4H), 3.77 (t, 2H), 3.52-3.50 (m, 2H), 3.42-3.36 (m, 5H), 3.33-3.29 (m, 2H), 3.18 (s, 3H), 3.00-2.94 (m, 2H), 2.24 (t, 2H), 2.03 (t, 2H), 1.87-1.73 (m, 2H), 1.69-1.67 (m, 2H), 1.64 (s, 12H), 1.55-1.36 (m, 4H), 1.35- 1.20 (m, 6H). $^{13}$C NMR (101 MHz, DMSO-$d_6$) δ 175.01, 174.67, 174.20, 172.17,157.80, 145.33, 142.89, 142.71, 141.34, 140.67, 128.74, 126.62, 125.97, 125.10, 122.72, 120.23, 112.00, 110.44, 71.68, 70.76, 70.25, 70.11, 67.80, 58.50, 52.74, 52.14, 49.23, 49.00, 44.44, 43.94, 40.65, 40.59, 38.69, 35.60, 32.16, 30.37, 29.28, 27.93, 27.67, 27.63, 27.19, 26.22, 25.40, 23.06. LC-MS(ESI) $[M+H]^+$ m/z found 1022.3, calcd for $C_{52}H_{72}N_5O_{14}S$ 1022.47.

### NMR and HPLC-MS characterization

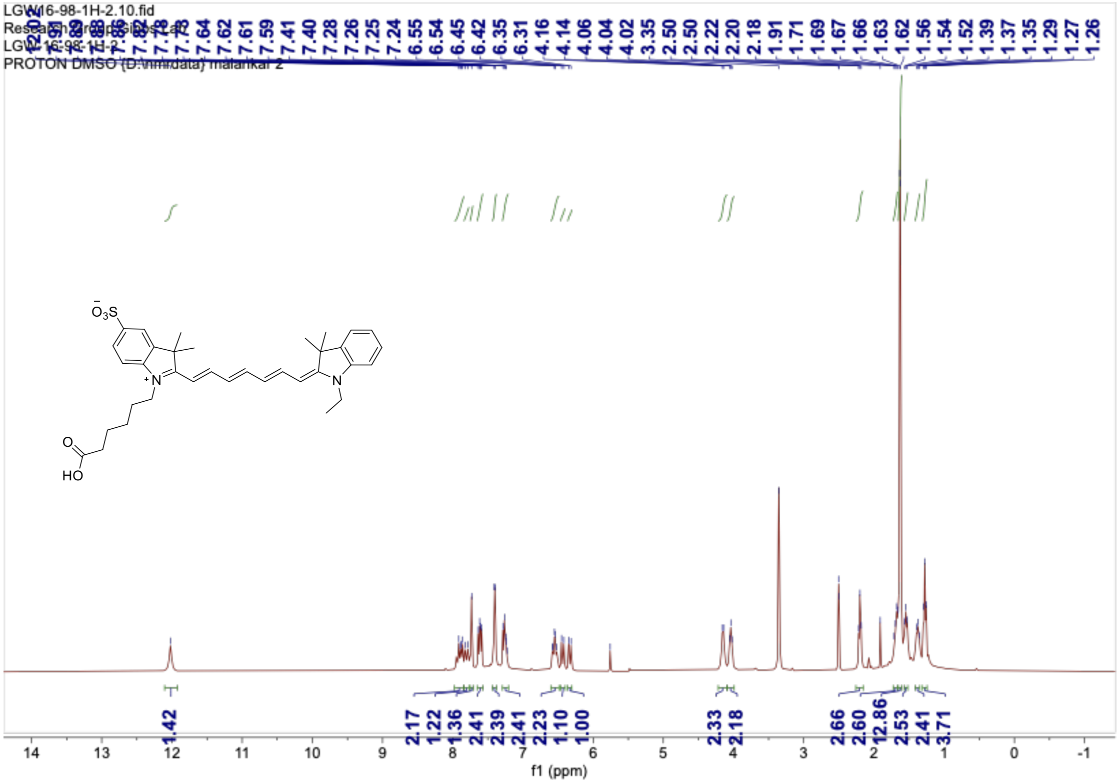

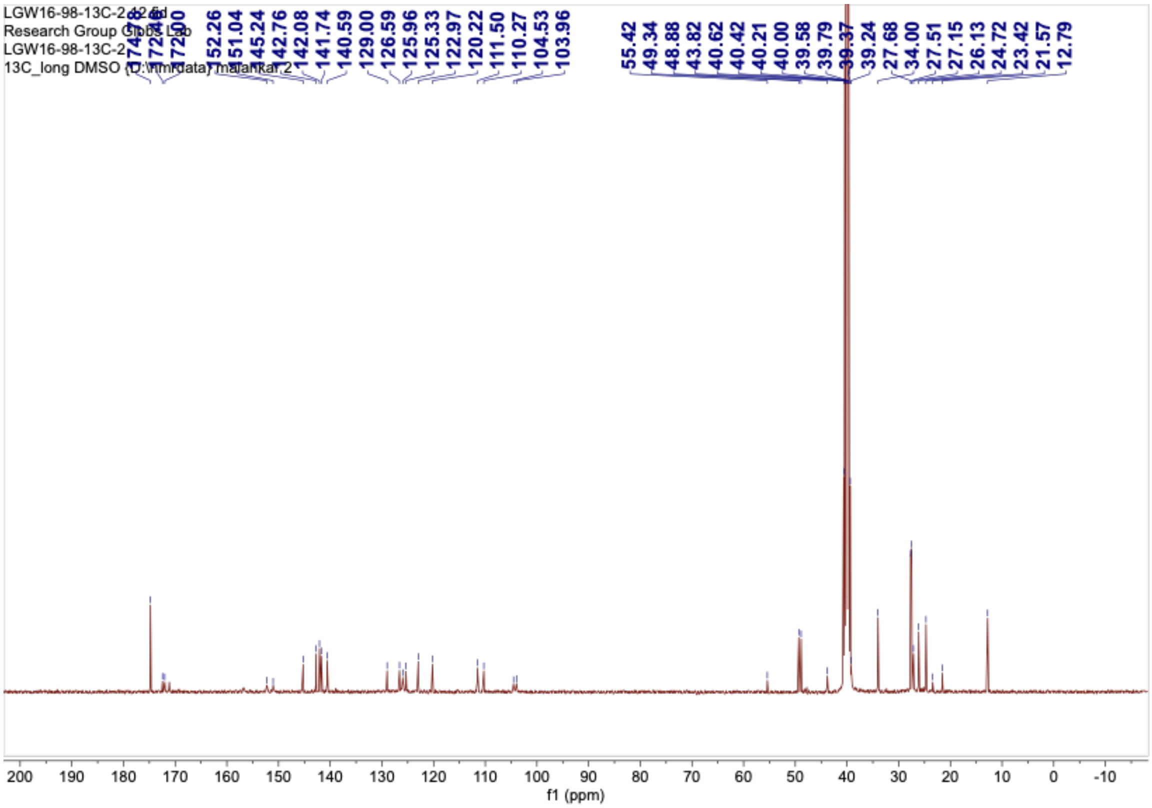

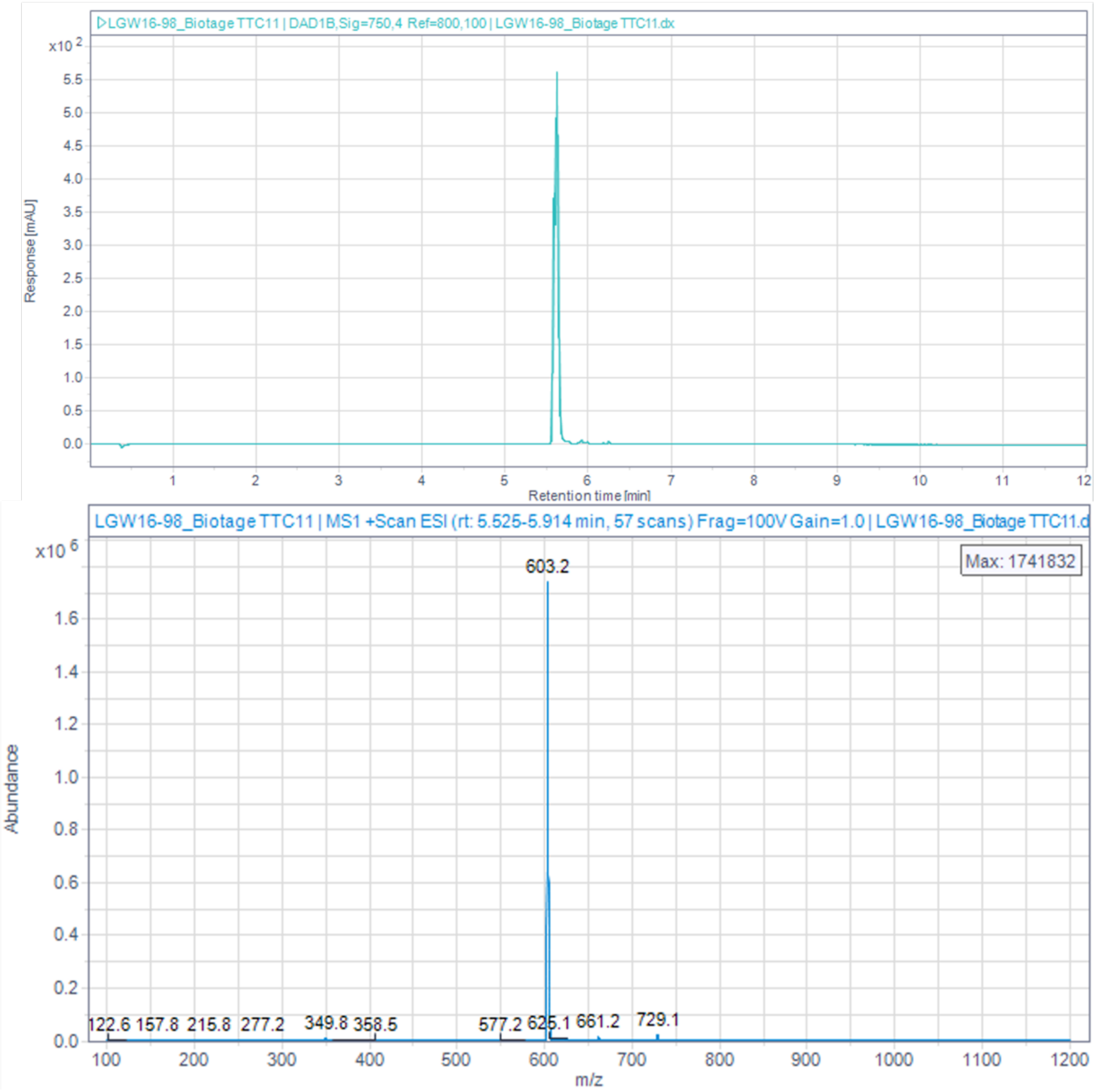


**Figure S3**. 1D NMR and LCMS characterization of intermediate **11**.

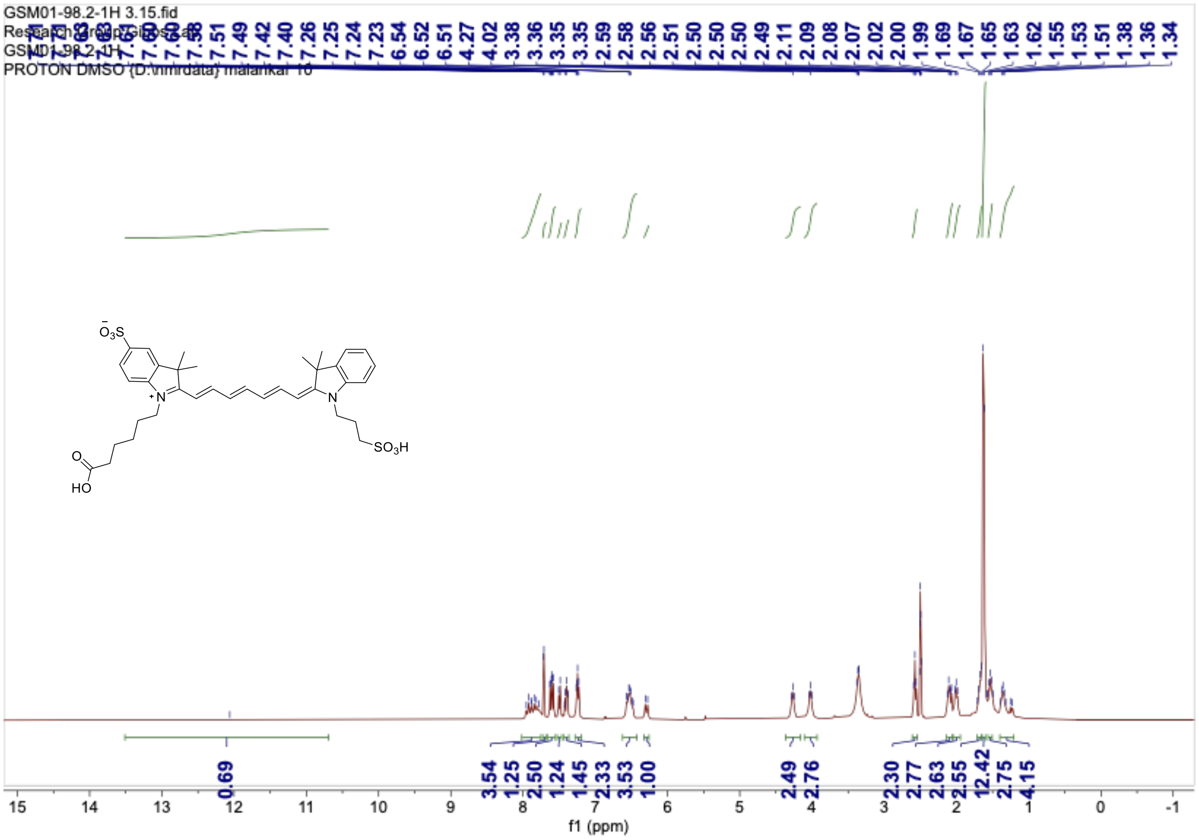
GSM01-98.2-1H 3.15.fid
PROTON DMSO
7.51 7.49 7.42 7.40 7.26 7.25 7.24 7.23 6.54 6.52 6.51 4.27 4.02 3.38 3.36 3.35 3.35 2.59 2.58 2.56 2.51 2.50 2.50 2.50 2.49 2.11 2.09 2.08 2.07 2.02 2.00 1.99 1.69 1.67 1.65 1.63 1.62 1.55 1.53 1.51 1.38 1.36 1.34
0.69 3.54 1.25 2.50 1.24 1.45 2.33 3.53 1.00 2.49 2.76 2.30 2.77 2.63 2.55 12.42 2.75 4.15
f1 (ppm)

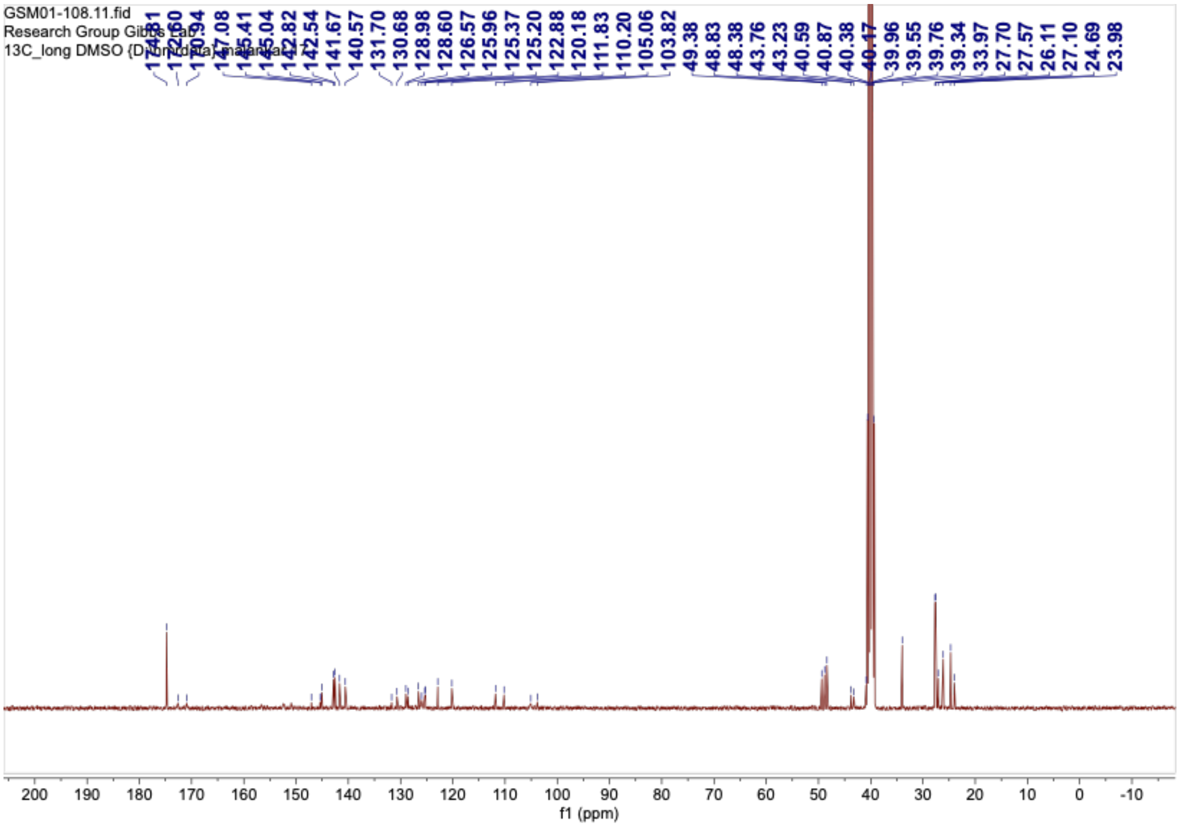
GSM01-108.11.fid
Research Group
13C_long DMSO
172.60 170.94 145.41 145.04 142.82 142.54 141.67 140.57 131.70 130.68 128.98 128.60 126.57 125.96 125.37 125.20 122.88 120.18 111.83 110.20 105.06 103.82 49.38 48.83 48.38 43.76 43.23 40.59 40.87 40.38 39.96 39.55 39.76 39.34 33.97 27.70 27.57 26.11 27.10 24.69 23.98
f1 (ppm)

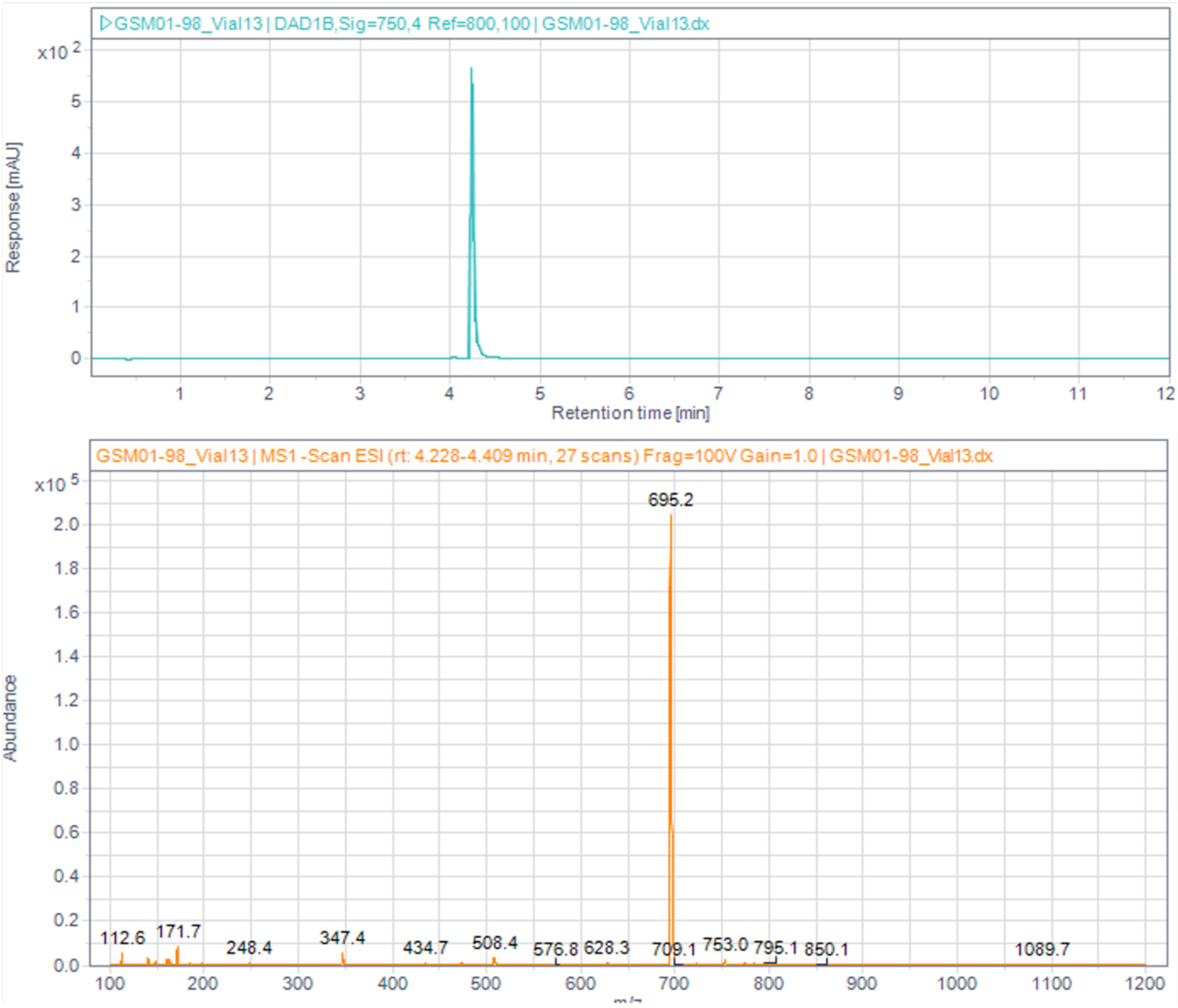


**Figure S4.** 1D NMR and LCMS characterization of intermediate **12**.

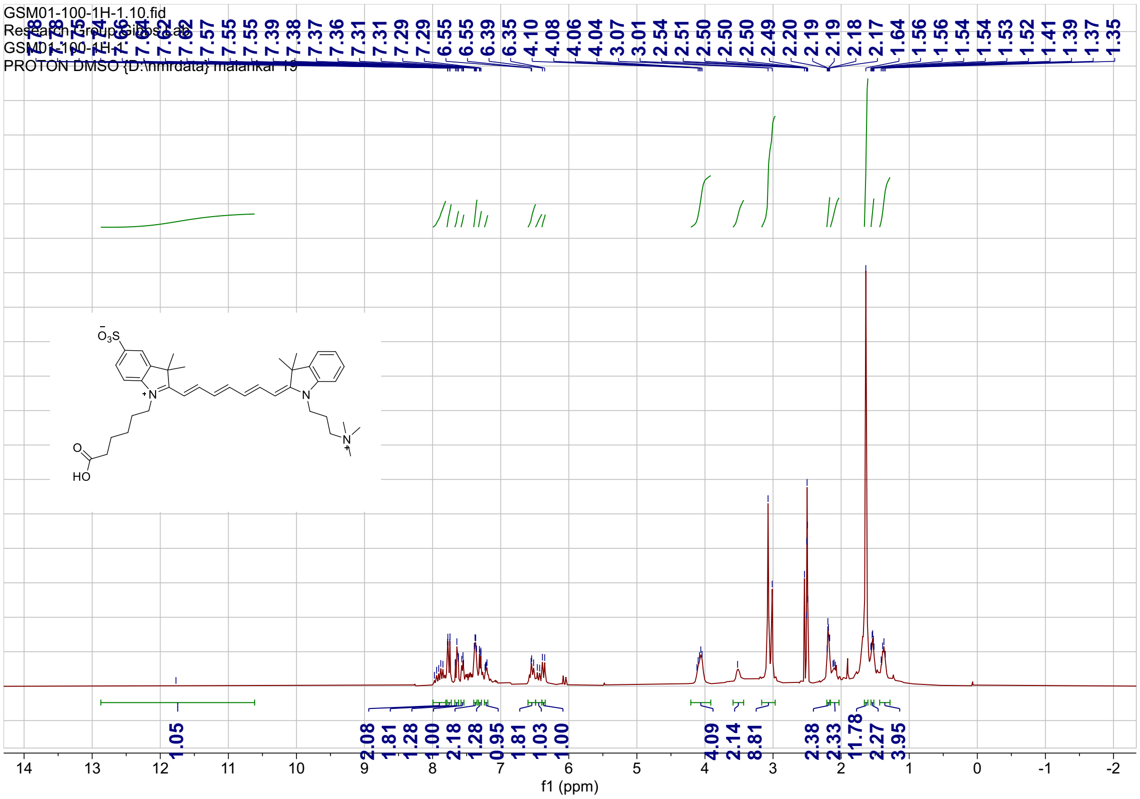
GSM01-100-1H-1.10.fid
PROTON DMSO {D:\nmrdata} malankar 19
f1 (ppm)

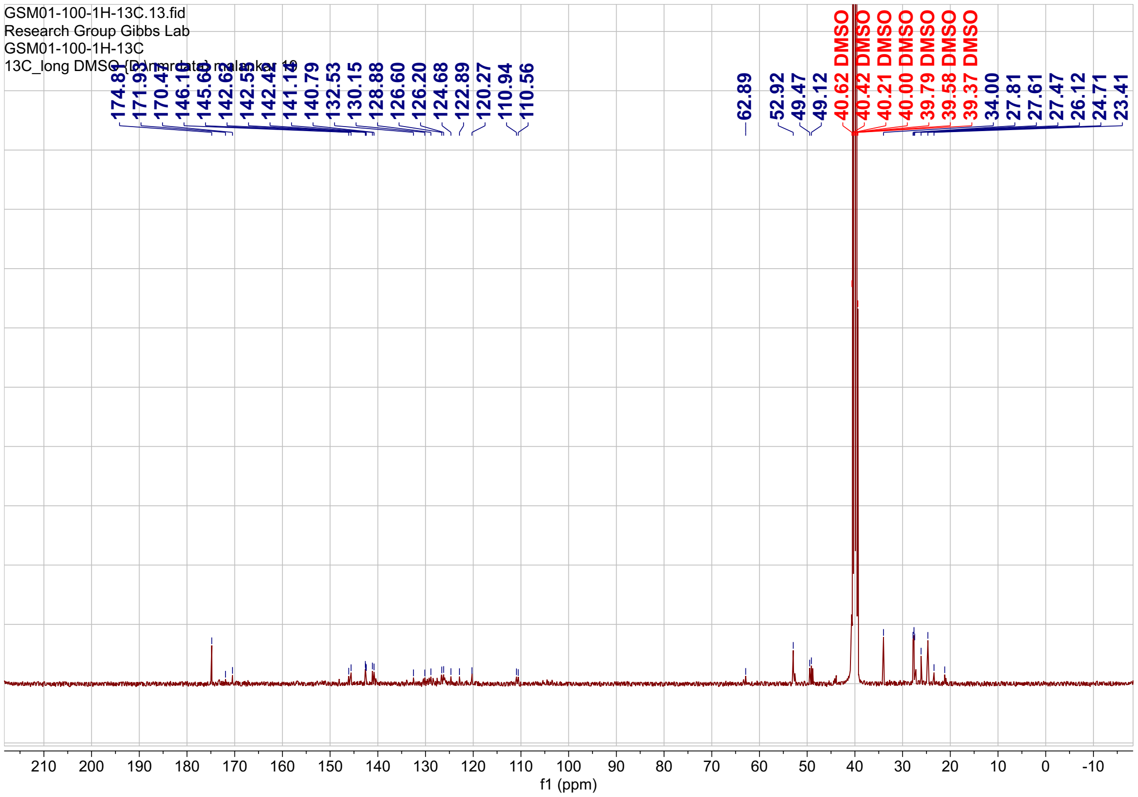
GSM01-100-1H-13C.13.fid
Research Group Gibbs Lab
GSM01-100-1H-13C
174.81
62.89
52.92
49.47
49.12
40.62 DMSO
40.42 DMSO
40.21 DMSO
40.00 DMSO
39.79 DMSO
39.58 DMSO
39.37 DMSO
34.00
27.81
27.61
27.47
26.12
24.71
23.41
f1 (ppm)

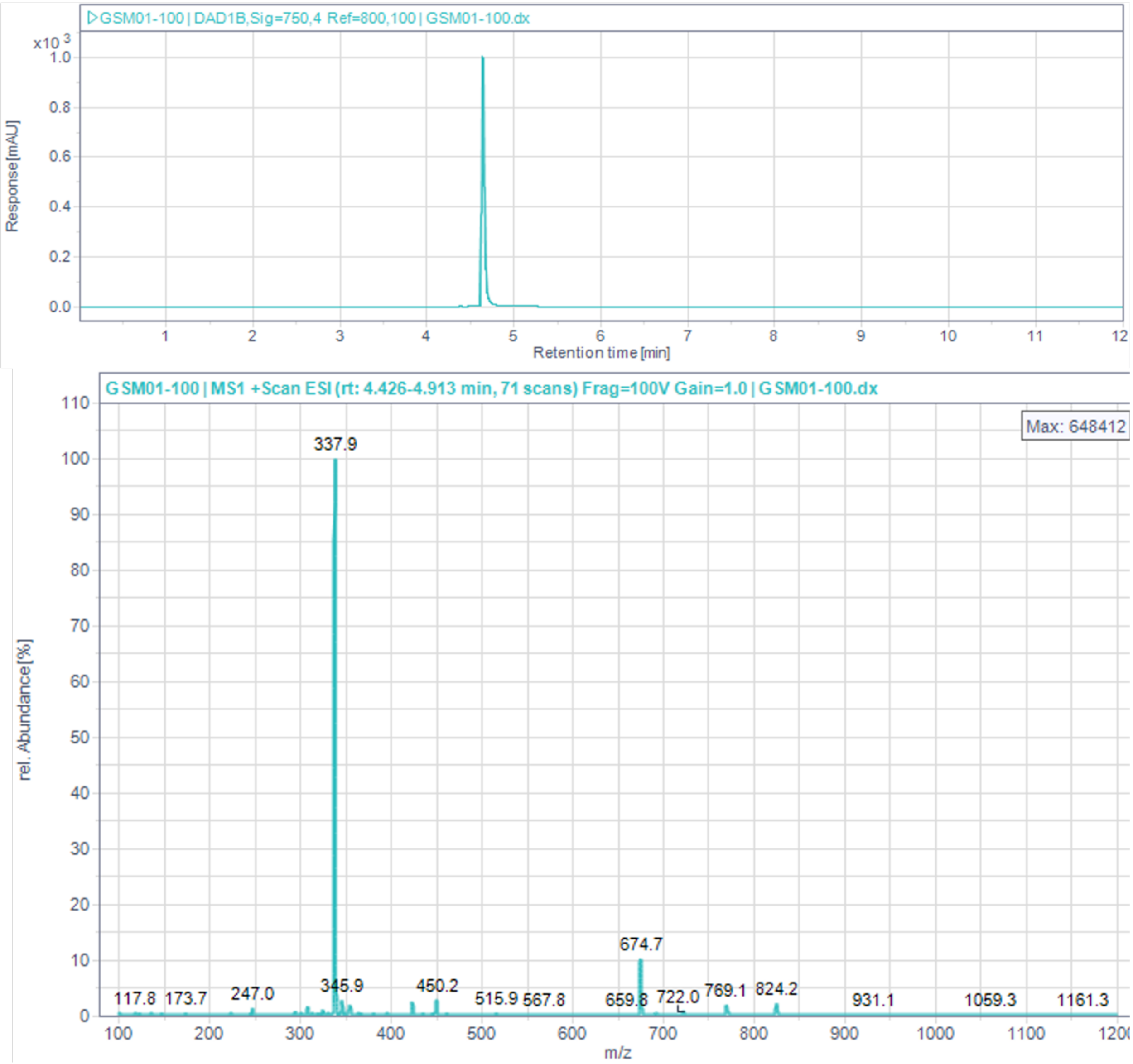


**Figure S5**. 1D NMR and LCMS characterization of intermediate **13**.

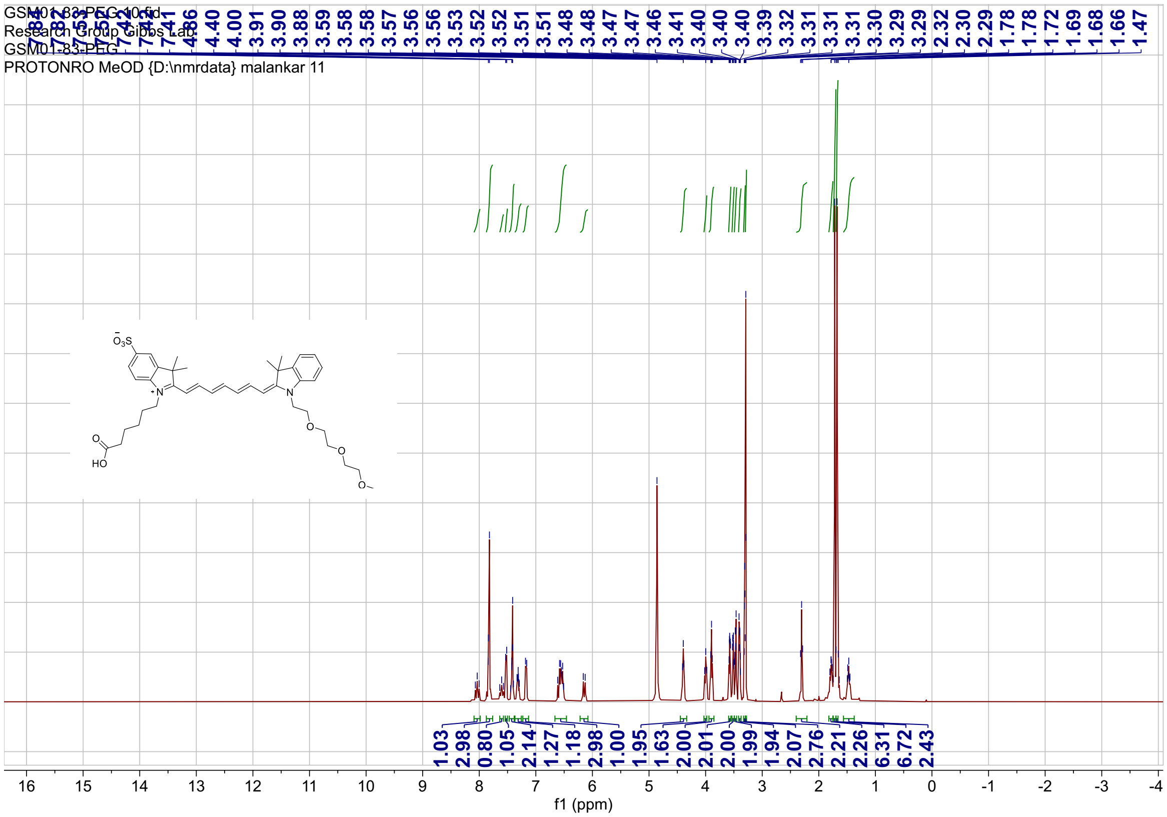
GSM01-83-PEG.10.fid
Research Group Gibbs Lab
GSM01-83-PEG
PROTONRO MeOD {D:\nmrdata} malankar 11
4.86 4.40 4.00 3.91 3.90 3.88 3.59 3.58 3.58 3.57 3.56 3.56 3.53 3.52 3.52 3.51 3.51 3.48 3.48 3.47 3.46 3.41 3.40 3.40 3.40 3.39 3.32 3.31 3.31 3.31 3.30 3.29 2.32 2.30 2.29 1.78 1.72 1.69 1.68 1.66 1.47
1.03 2.98 0.80 1.05 2.14 1.27 1.18 2.98 1.00 1.95 1.63 2.00 2.01 2.00 1.99 1.94 2.07 2.76 2.21 2.26 6.31 6.72 2.43
f1 (ppm)

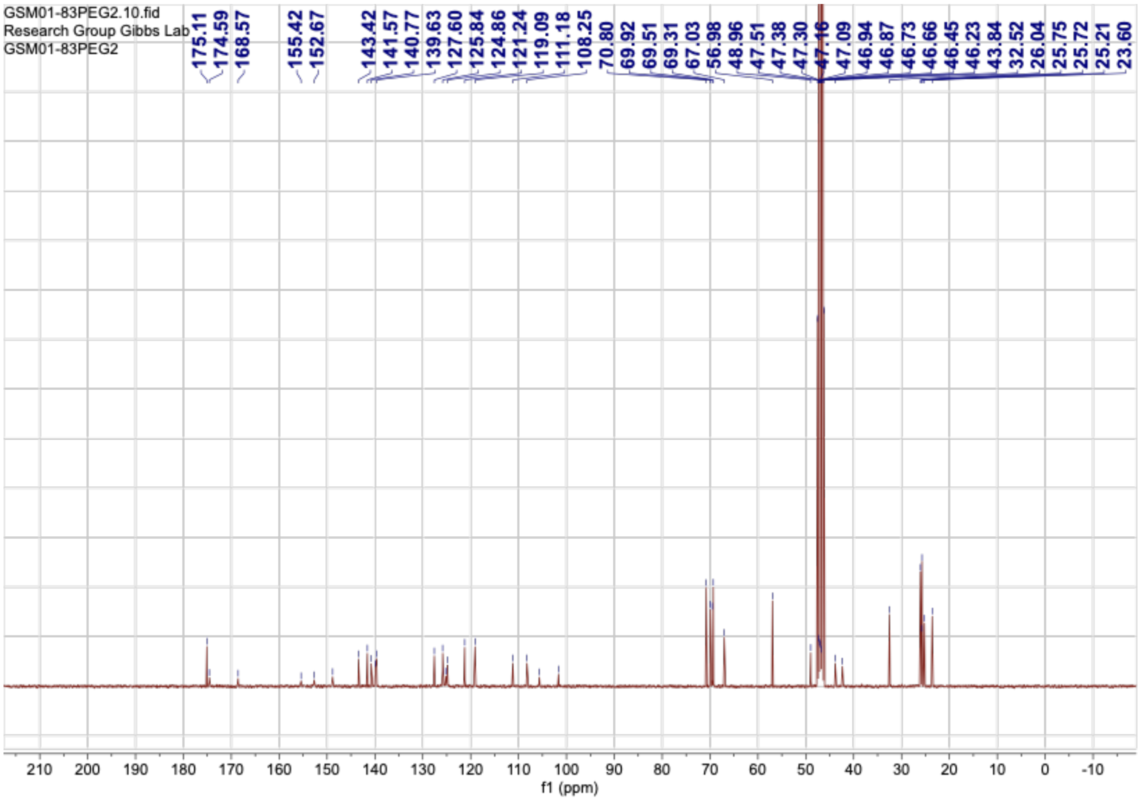
GSM01-83PEG2.10.fid
Research Group Gibbs Lab
GSM01-83PEG2
175.11 174.59 168.57 155.42 152.67 143.42 141.57 140.77 139.63 127.60 125.84 124.86 121.24 119.09 111.18 108.25 70.80 69.92 69.51 69.31 67.03 56.98 48.96 47.51 47.38 47.30 47.16 47.09 46.94 46.87 46.73 46.66 46.45 46.23 43.84 32.52 26.04 25.75 25.72 25.21 23.60
f1 (ppm)

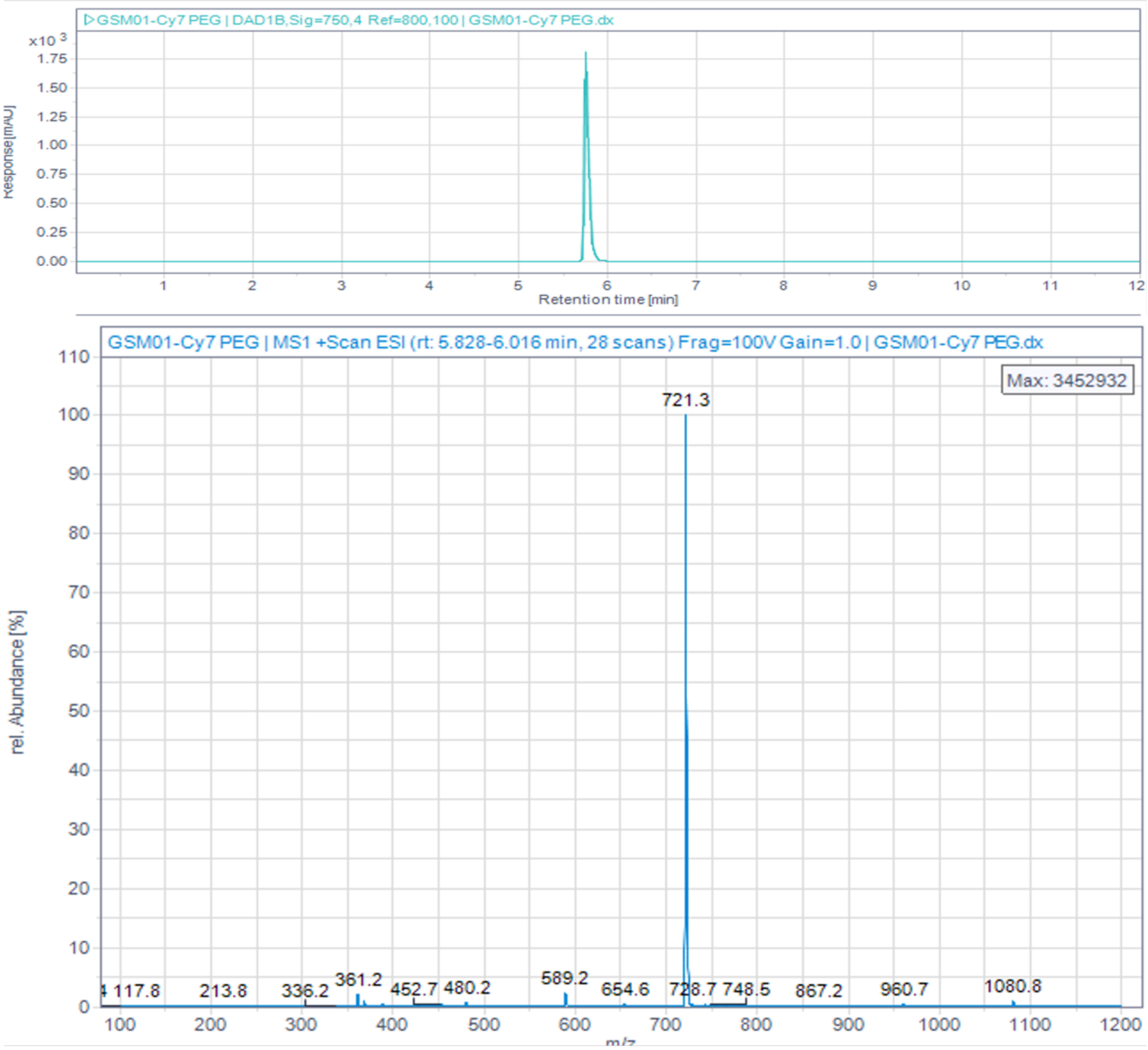


**Figure S6**. 1D NMR and LCMS characterization of intermediate **14**.

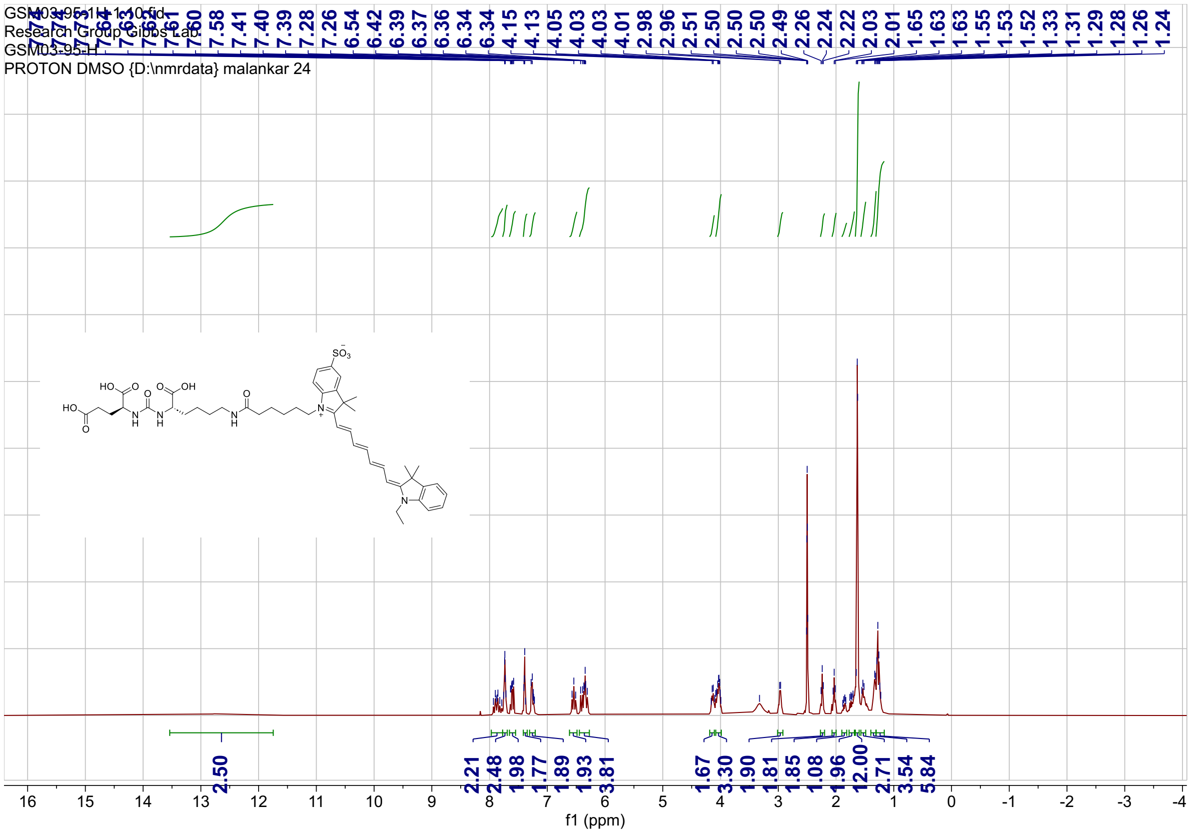
GSM03-95-1H-1-10.fid
Research Group Gibbs Lab
GSM03-95-H
PROTON DMSO {D:\nmrdata} malankar 24
7.60 7.58 7.41 7.40 7.39 7.28 7.26 6.54 6.42 6.39 6.37 6.36 6.34 6.34 4.15 4.13 4.05 4.03 4.03 4.01 2.98 2.96 2.51 2.50 2.50 2.50 2.49 2.26 2.24 2.22 2.03 2.01 1.65 1.63 1.63 1.55 1.53 1.52 1.33 1.31 1.29 1.28 1.26 1.24
2.50 2.21 2.48 1.98 1.77 1.89 1.93 3.81 1.67 3.30 1.90 1.81 1.85 1.08 1.96 12.00 2.71 3.54 5.84
16 15 14 13 12 11 10 9 8 7 6 5 4 3 2 1 0 -1 -2 -3 -4
f1 (ppm)

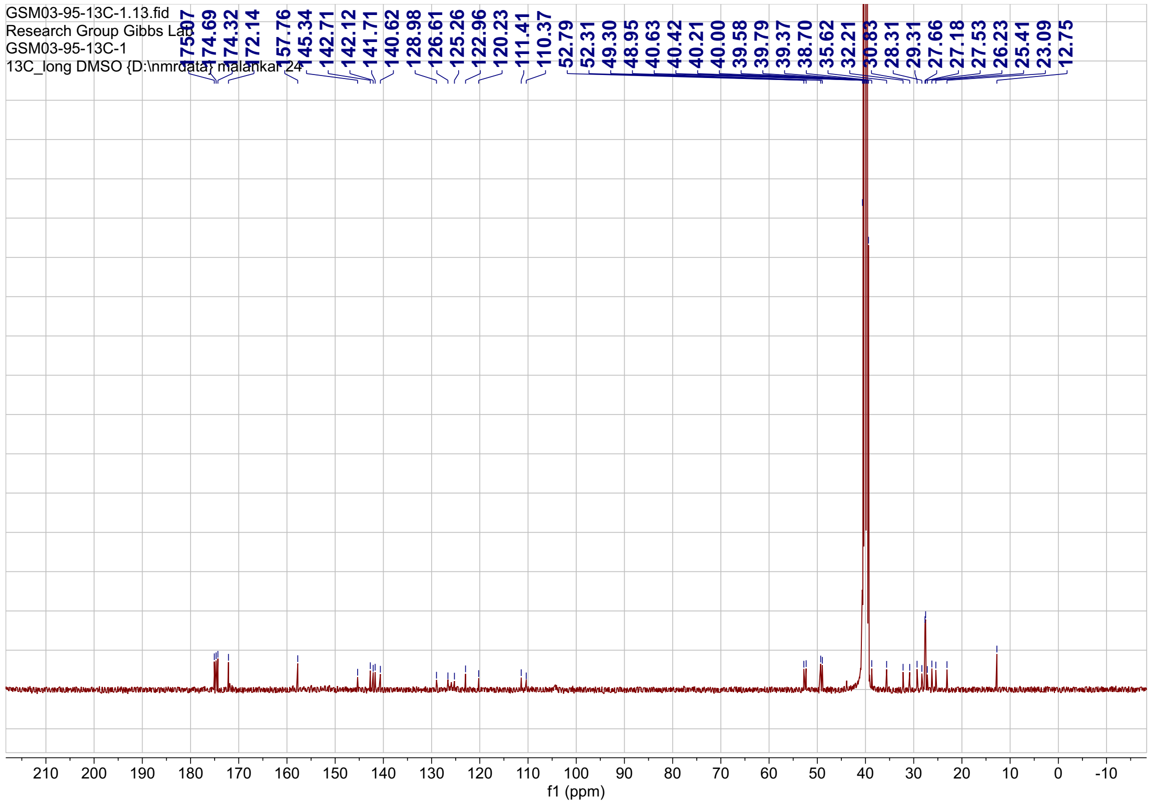
GSM03-95-13C-1.13.fid
Research Group Gibbs Lab
GSM03-95-13C-1
13C_long DMSO {D:\nmrdata} malankar 24
175.07 174.69 174.32 172.14 157.76 145.34 142.71 142.12 141.71 140.62 128.98 126.61 125.26 122.96 120.23 111.41 110.37 52.79 52.31 49.30 48.95 40.63 40.42 40.21 40.00 39.58 39.79 39.37 38.70 35.62 32.21 30.83 28.31 29.31 27.66 27.18 27.53 26.23 25.41 23.09 12.75
210 200 190 180 170 160 150 140 130 120 110 100 90 80 70 60 50 40 30 20 10 0 -10
f1 (ppm)

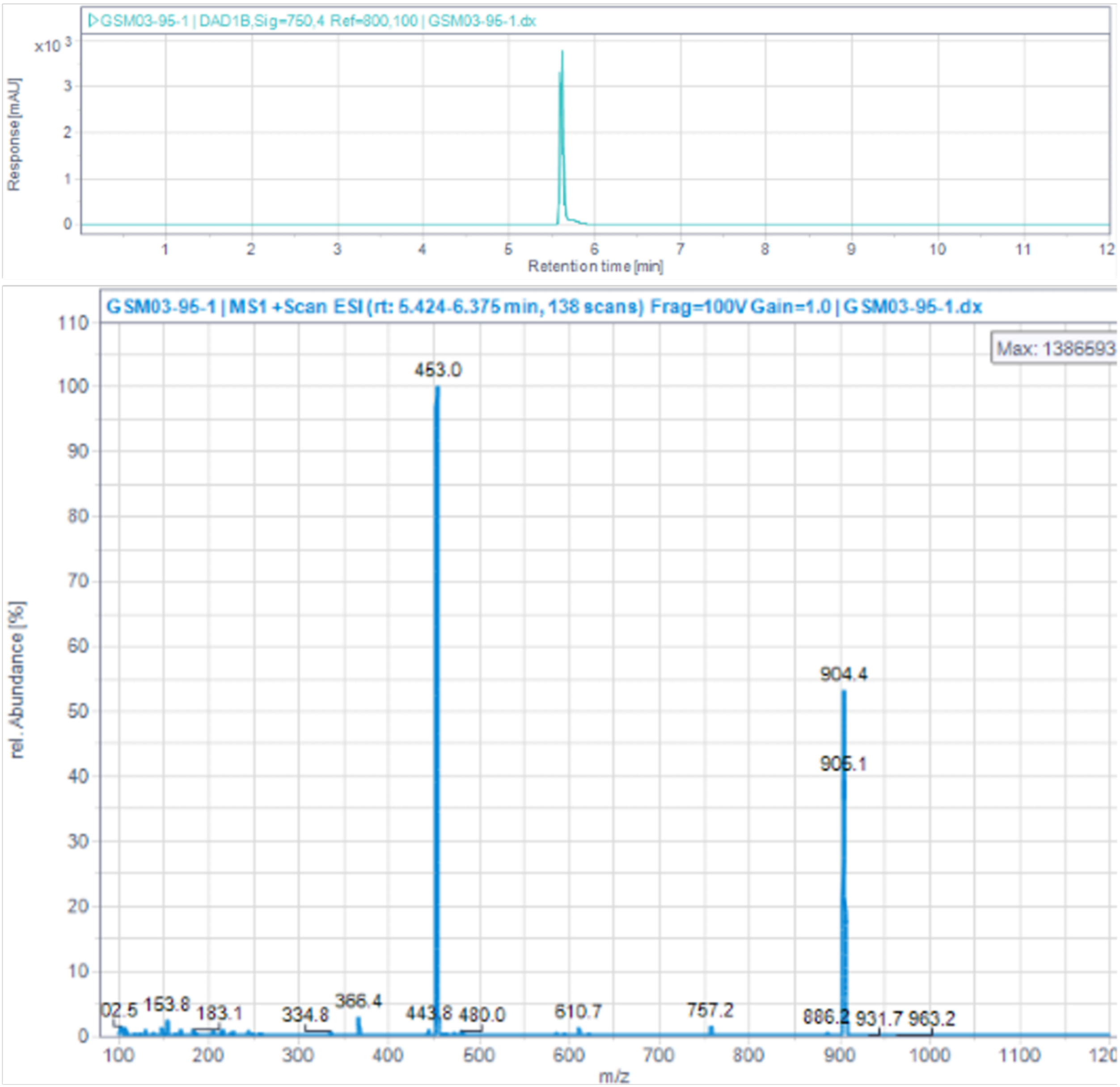


**Figure S7**. 1D NMR and LCMS characterization of PSMA-1 (Compound **16**).

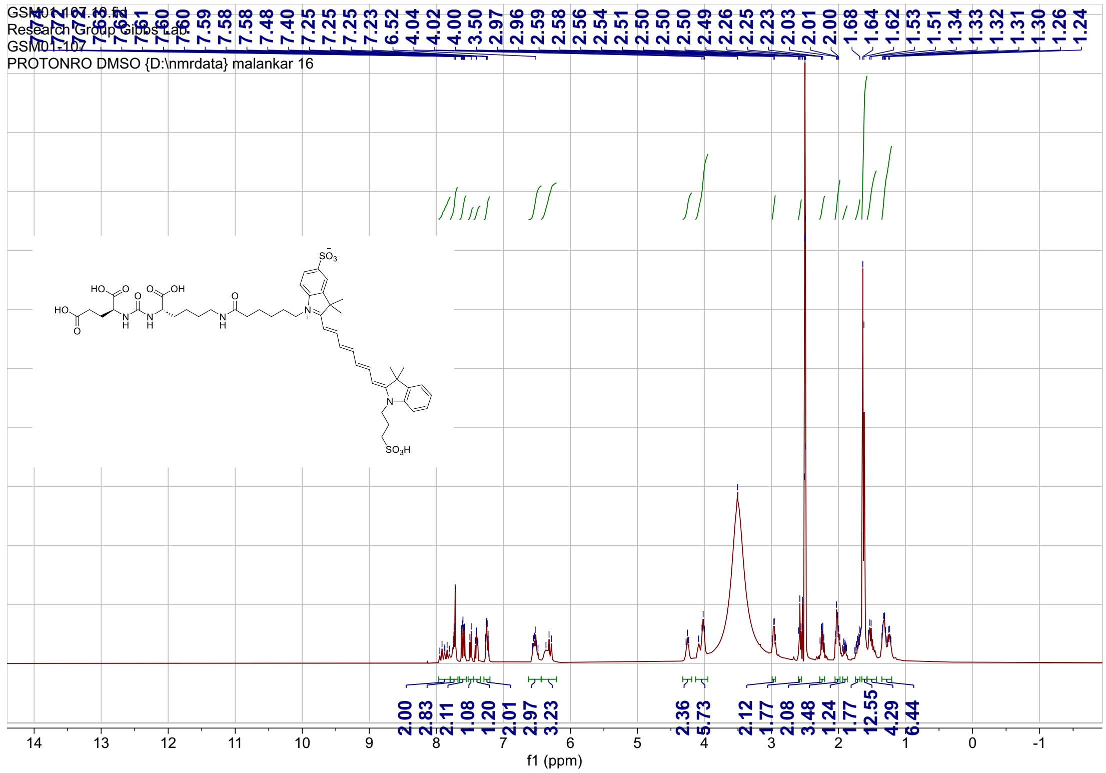
Research Group Gibbs Lab
GSM01-107
PROTONRO DMSO {D:\nmrdata} malankar 16
f1 (ppm)

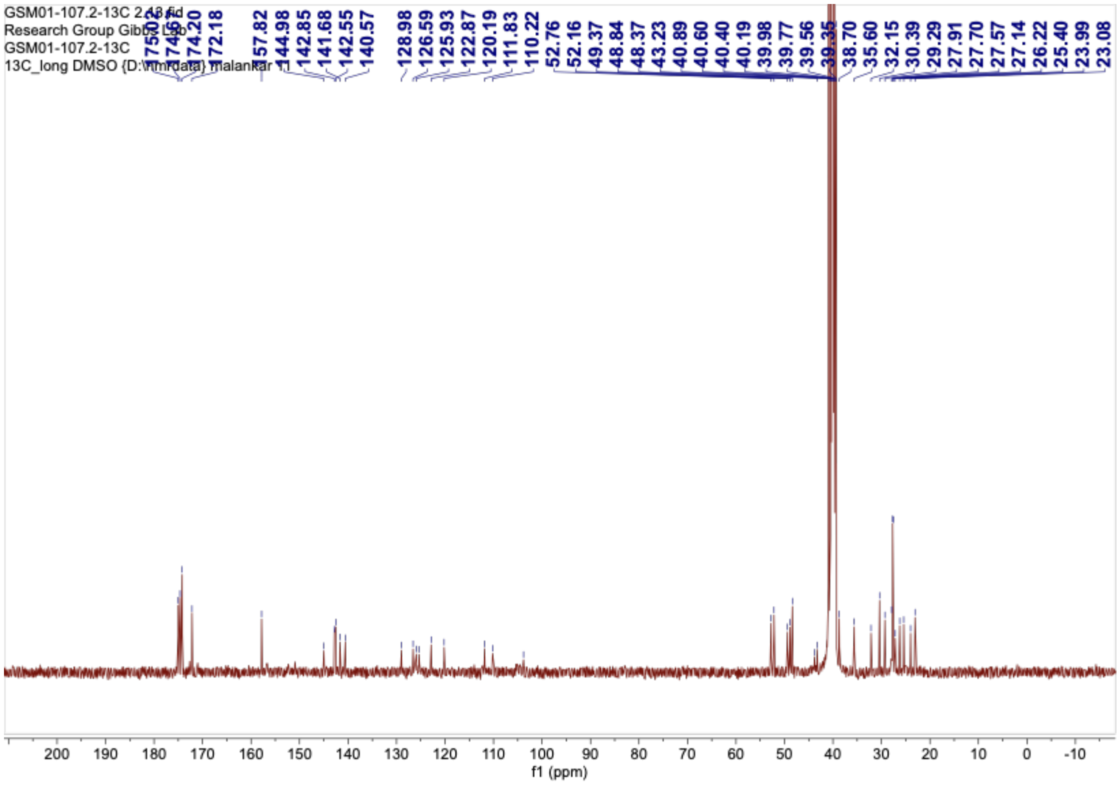
GSM01-107.2-13C
Research Group Gibbs Lab
GSM01-107.2-13C
f1 (ppm)

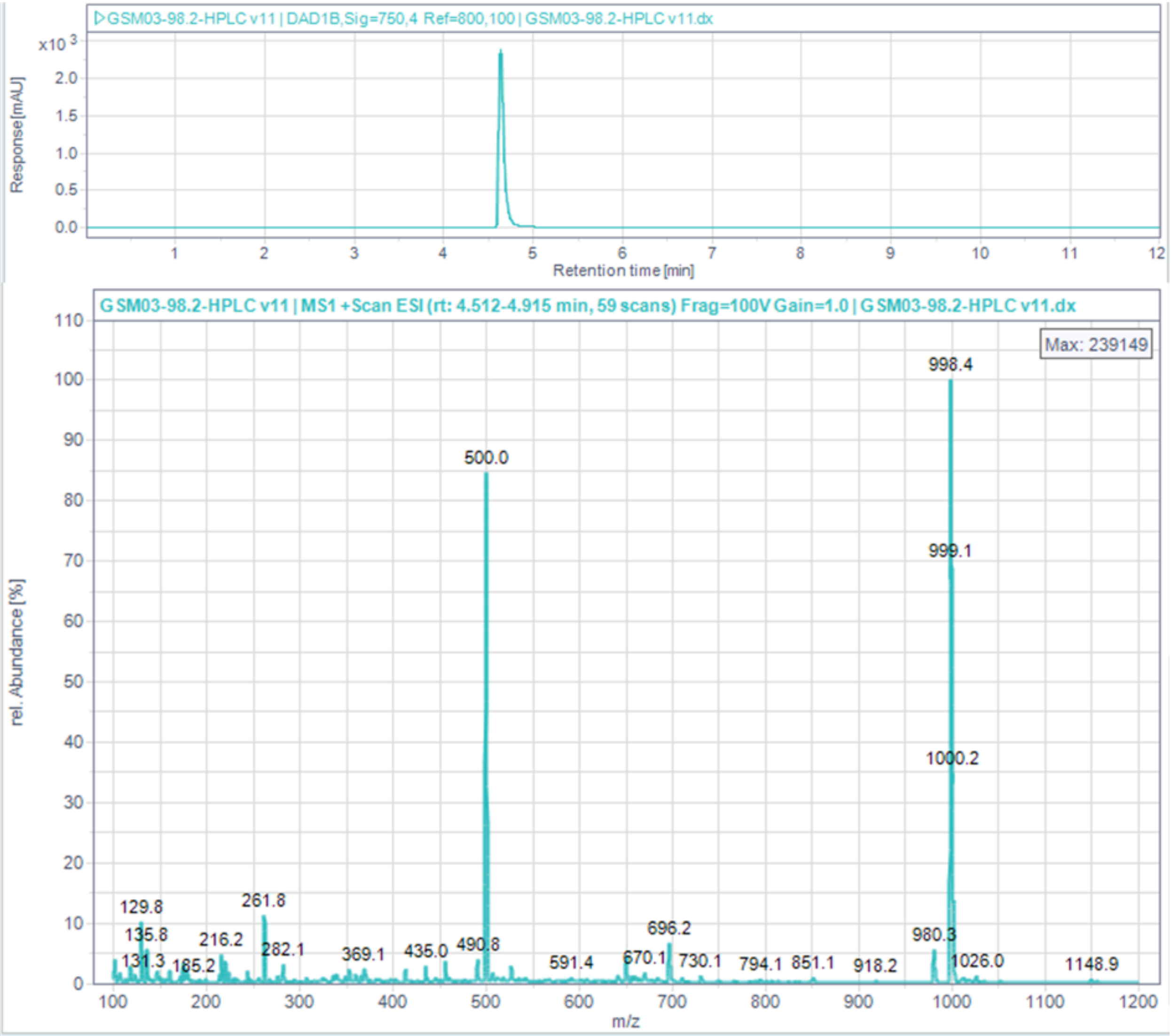


**Figure S8**. 1D NMR and LCMS characterization of PSMA-2 (Compound **17**).

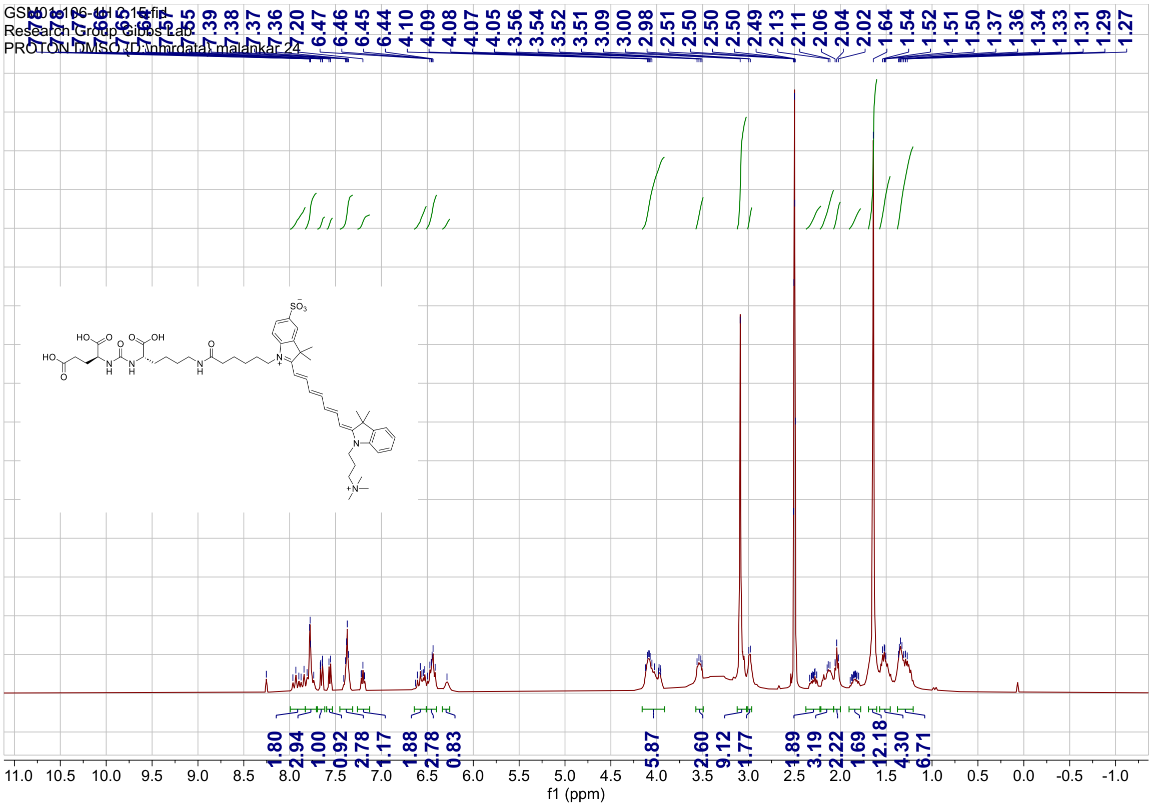
Research Group Gibbs Lab
PROTON DMSO {D:\nmrdata} malankar 24
f1 (ppm)

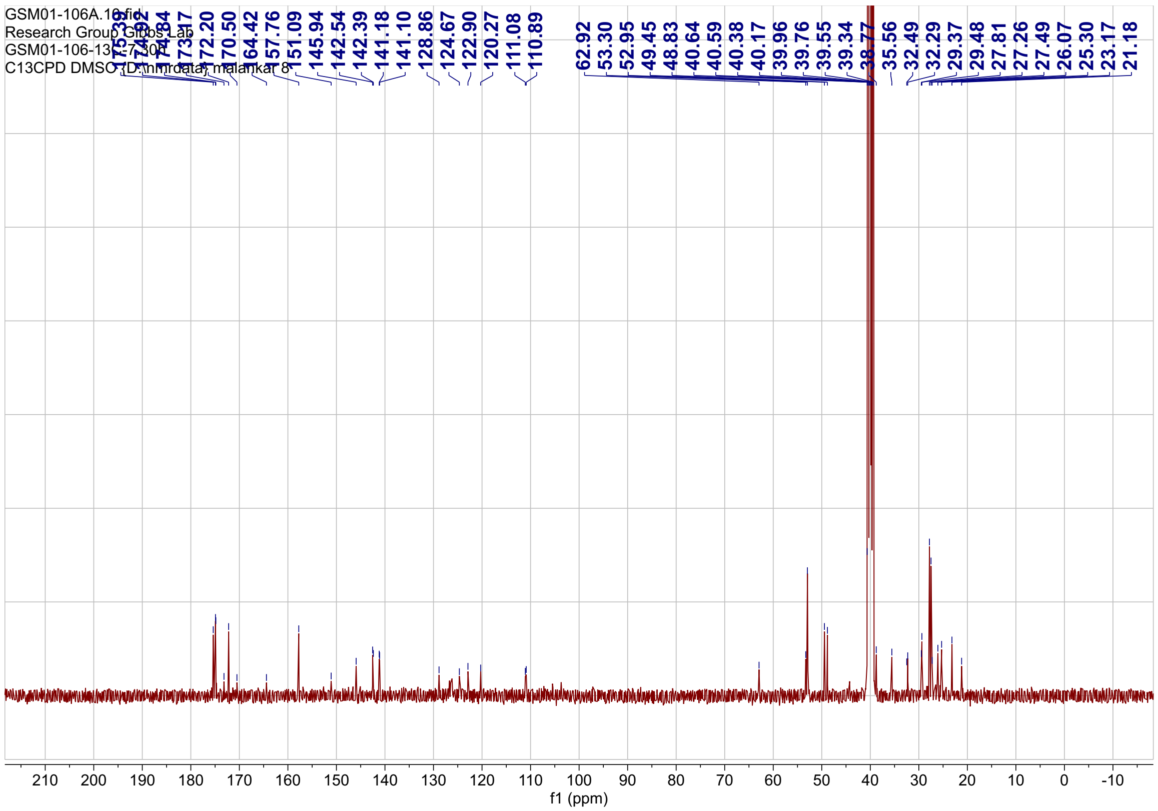
GSM01-106A.10.fid
Research Group Gibbs Lab
C13CPD DMSO {D:\nmrdata} malankar 8
f1 (ppm)

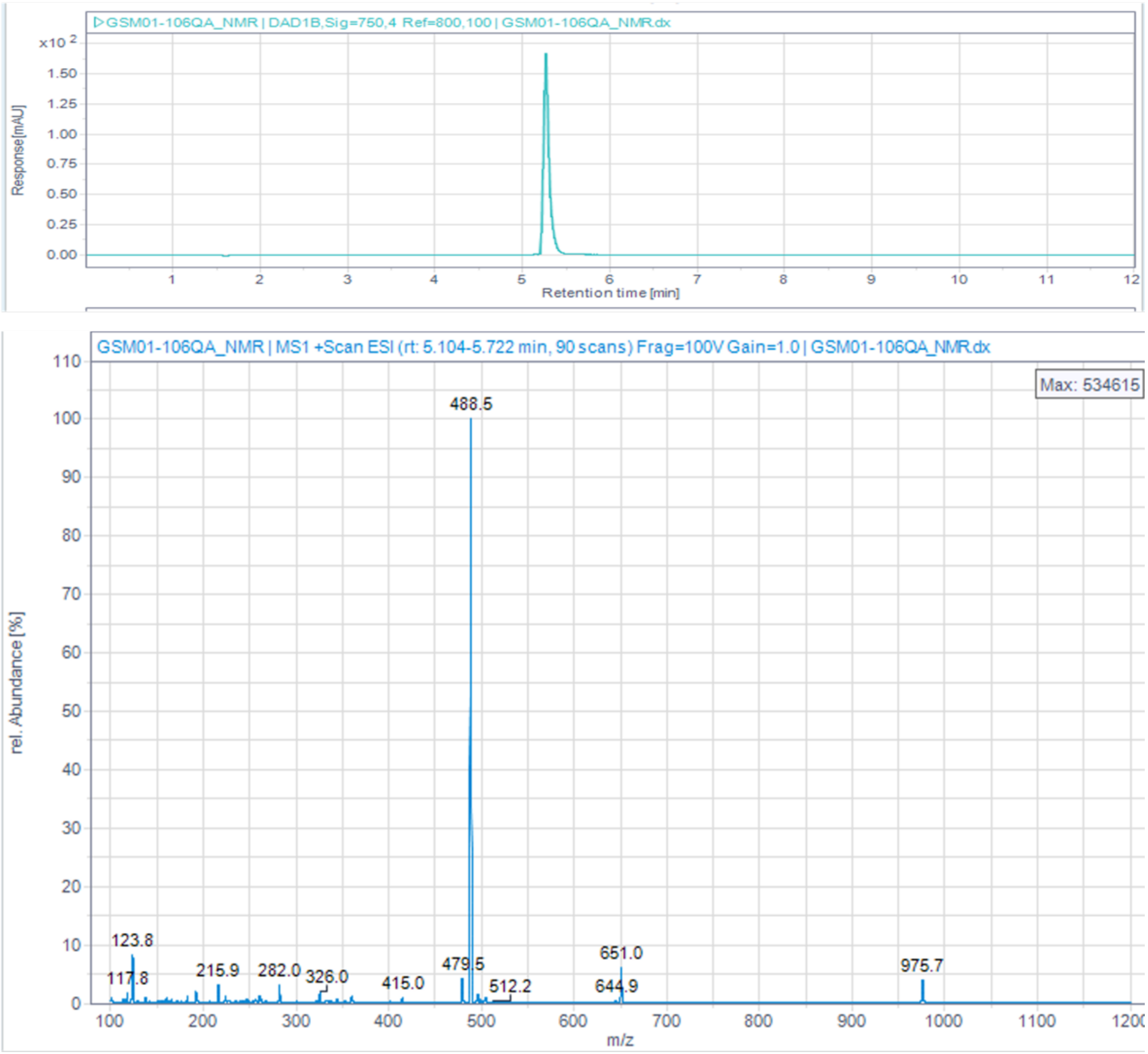


**Figure S9**. 1D NMR and LCMS characterization of PSMA-3 (Compound **18**).

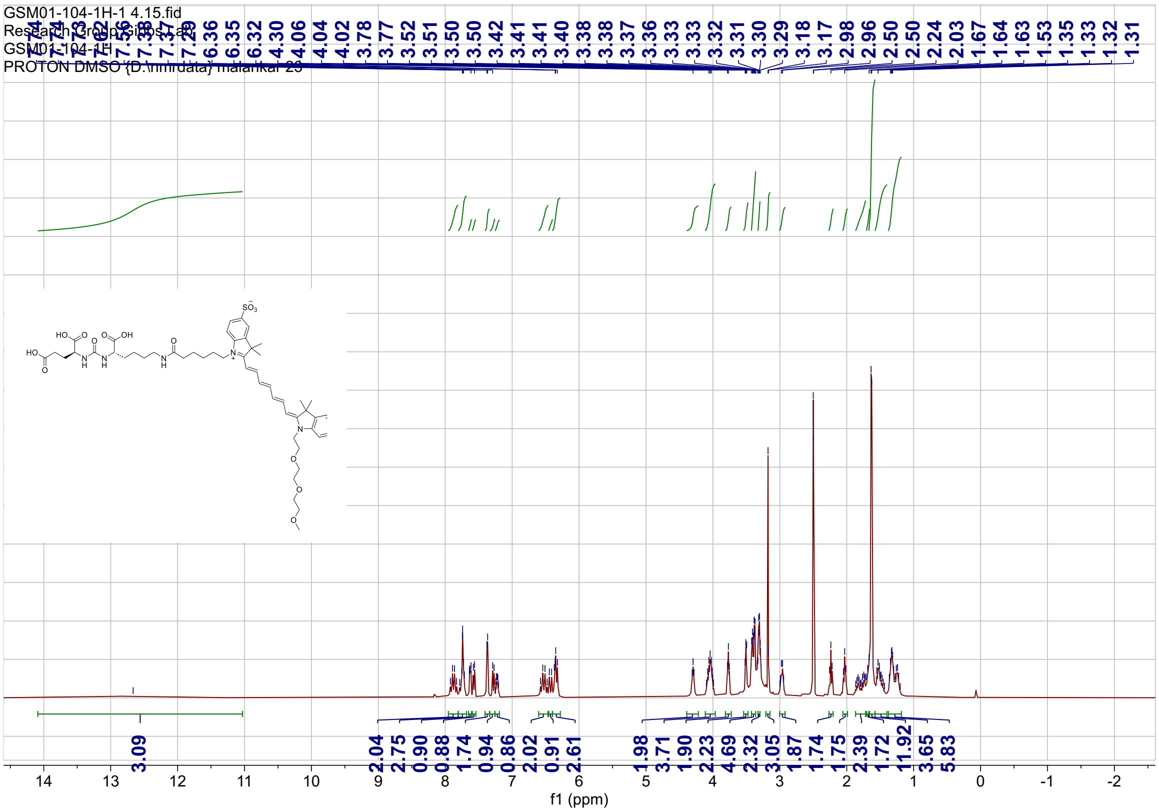
GSM01-104-1H-1 4.15.fid
GSM01-104-1H
PROTON DMSO {D:\nmrdata} malankar 23
6.36 6.35 6.32 4.30 4.06 4.04 4.02 3.78 3.77 3.52 3.51 3.50 3.50 3.42 3.41 3.40 3.38 3.38 3.37 3.36 3.33 3.33 3.32 3.31 3.30 3.29 3.18 3.17 2.98 2.96 2.50 2.50 2.24 2.03 1.67 1.64 1.63 1.53 1.35 1.33 1.32 1.31
3.09
2.04 2.75 0.90 0.88 1.74 0.94 0.86 2.02 0.91 2.61 1.98 3.71 1.90 2.23 4.69 2.32 3.05 1.87 1.74 1.75 2.39 1.72 11.92 3.65 5.83
f1 (ppm)

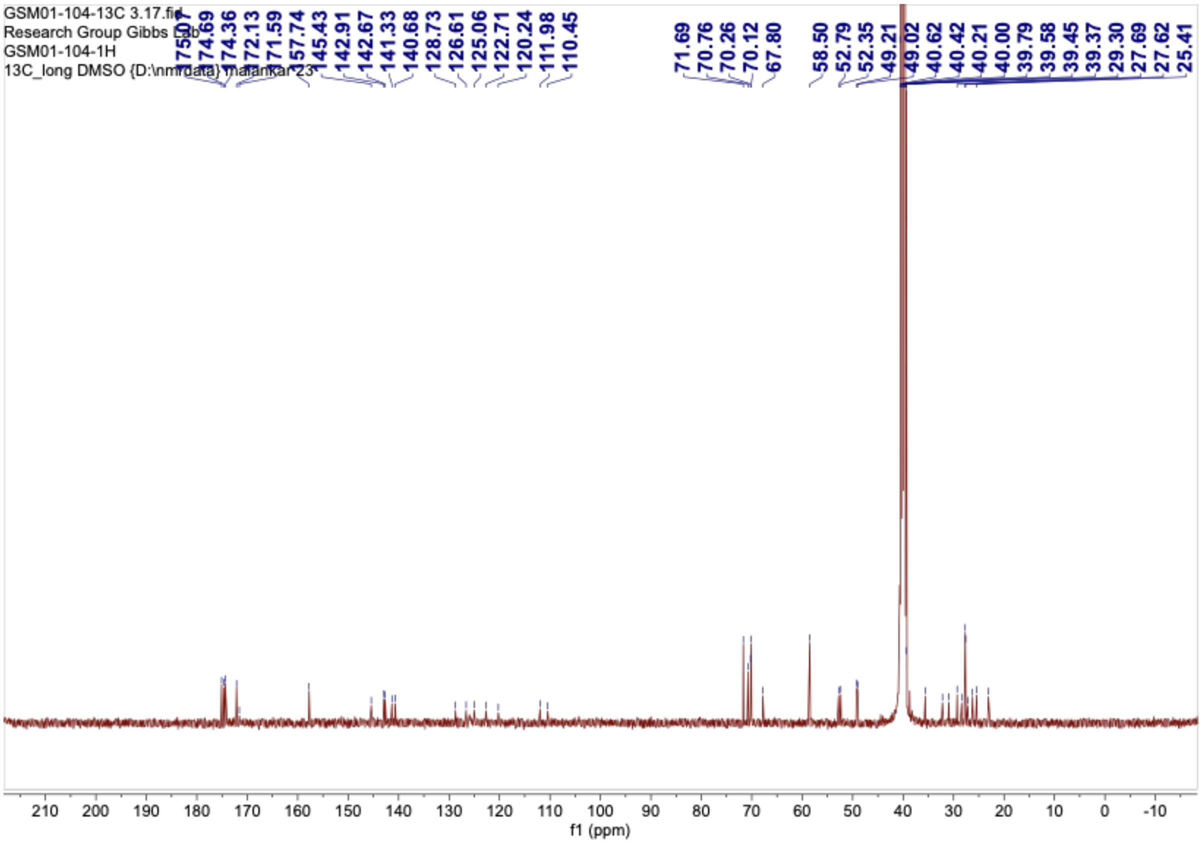
GSM01-104-13C 3.17.fid
Research Group Gibbs Lab
GSM01-104-1H
13C_long DMSO {D:\nmrdata} malankar 23
174.69 174.36 172.13 171.59 157.74 145.43 142.91 142.67 141.33 140.68 128.73 126.61 125.06 122.71 120.24 111.98 110.45 71.69 70.76 70.26 70.12 67.80 58.50 52.79 52.35 49.21 49.02 40.62 40.42 40.21 40.00 39.79 39.58 39.45 39.37 29.30 27.69 27.62 25.41
f1 (ppm)

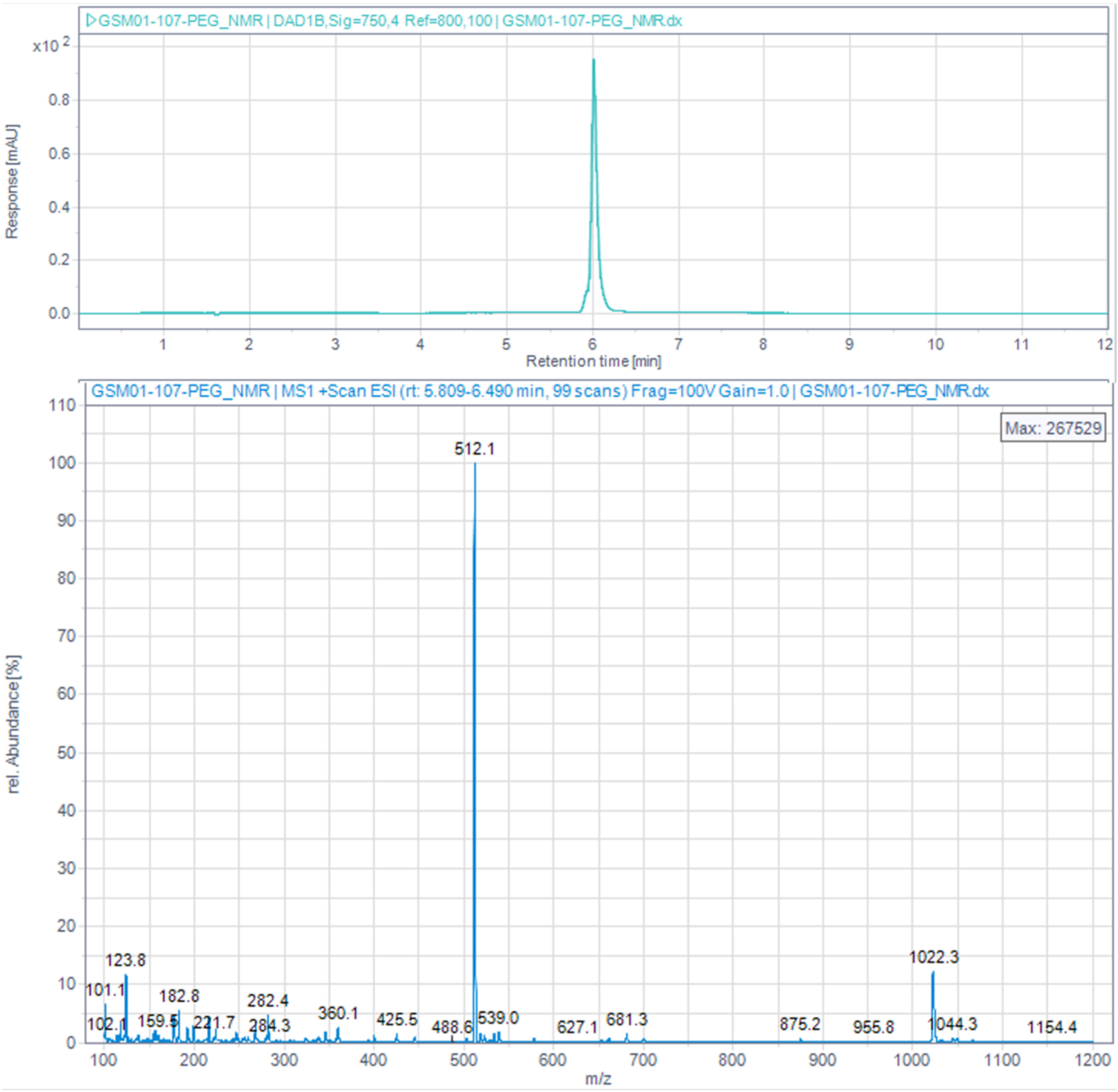


**Figure S10**. 1D NMR and LCMS characterization of PSMA-4 (Compound **19**).
